# The substrate-dependent stereoselectivity of the multicopper oxidase (MCO)-catalyzed oxidative coupling in the biosynthesis of the bisnaphthopyrone viriditoxin

**DOI:** 10.1101/846196

**Authors:** Jinyu Hu, Gavin R. Flematti, Yit-Heng Chooi

## Abstract

VdtB, the multiple-copper oxidase (MCO) from the bisnaphthopyrone (*M*)-viriditoxin biosynthetic pathway in *Paecilomyces variotii*, was shown to catalyze regioselective 6,6′-coupling of semi-viriditoxin (**1**). The stereoselectivity of the oxidative coupling reaction for the production of the atropisomer (*M*)-viriditoxin, however, was controlled by VdtD, a non-catalytic dirigent protein from the pathway. In this work, VdtB either alone or together with VdtD were investigated for its stereoselective control upon coupling of other monomeric naphthopyrone derivatives from the pathway with different minor structural variations in terms of presence/absence of *O*-methylation at C7-position and C3-C4 *Δ^2^* double bond on the pyrone ring, and the different side-chain modifications. We showed that VdtB could favour either *M*- or *P*-form coupling in a substrate-dependent manner. For some substrates, VdtB could catalyze oxidative coupling in an enantiomerically selective manner. The efficiency of the VdtD in exerting stereoselective control of the oxidative coupling reaction also varies between substrates. The results point to a model whereby VdtB and VdtD form a VdtB-ligand-VdtD complex in which the stereochemical outcome of the coupling reaction depends on how the substrate interacts with both proteins, based on the substrate structure. Our findings contributed to a more comprehensive understanding of dirigent protein-mediated MCO-catalyzed stereoselective oxidative coupling reactions in fungi.

## Main Text

Axially chirality in biaryl nature products has been gained growing interest during the past decade, both as a challenge in total synthetic point of view for controlling of the regio- and stereo-chemistry,^[1]^ and for its potential importance in pharmaceutical and medicinal chemistry.^[2]^ Significant efforts have been taken to investigate the stringent regio-and stereo-selective control in natural oxidative phenol coupling mechanism. In higher plants, the regio- and stereo-selectivity was conducted by the corporation of laccases/multicopper oxidases (LMCOs) and noncatalytic auxiliary proteins known as dirigent proteins.^[3]^ In *Streptomyces*, such selectivity was often attributed to members from cytochrome P450 enzymes (P450s).^[4]^ In fungi, members of P450s have been characterized as regio- and stereo-selective coupling enzyme,^[5]^ while recent biosynthetic studies featured the discovery of MCOs that, either alone or together with accessory proteins, are also capable of catalyzing such regio- and stereo-selective coupling reactions (Figure 1A).^[6]^

**Figure 1.**
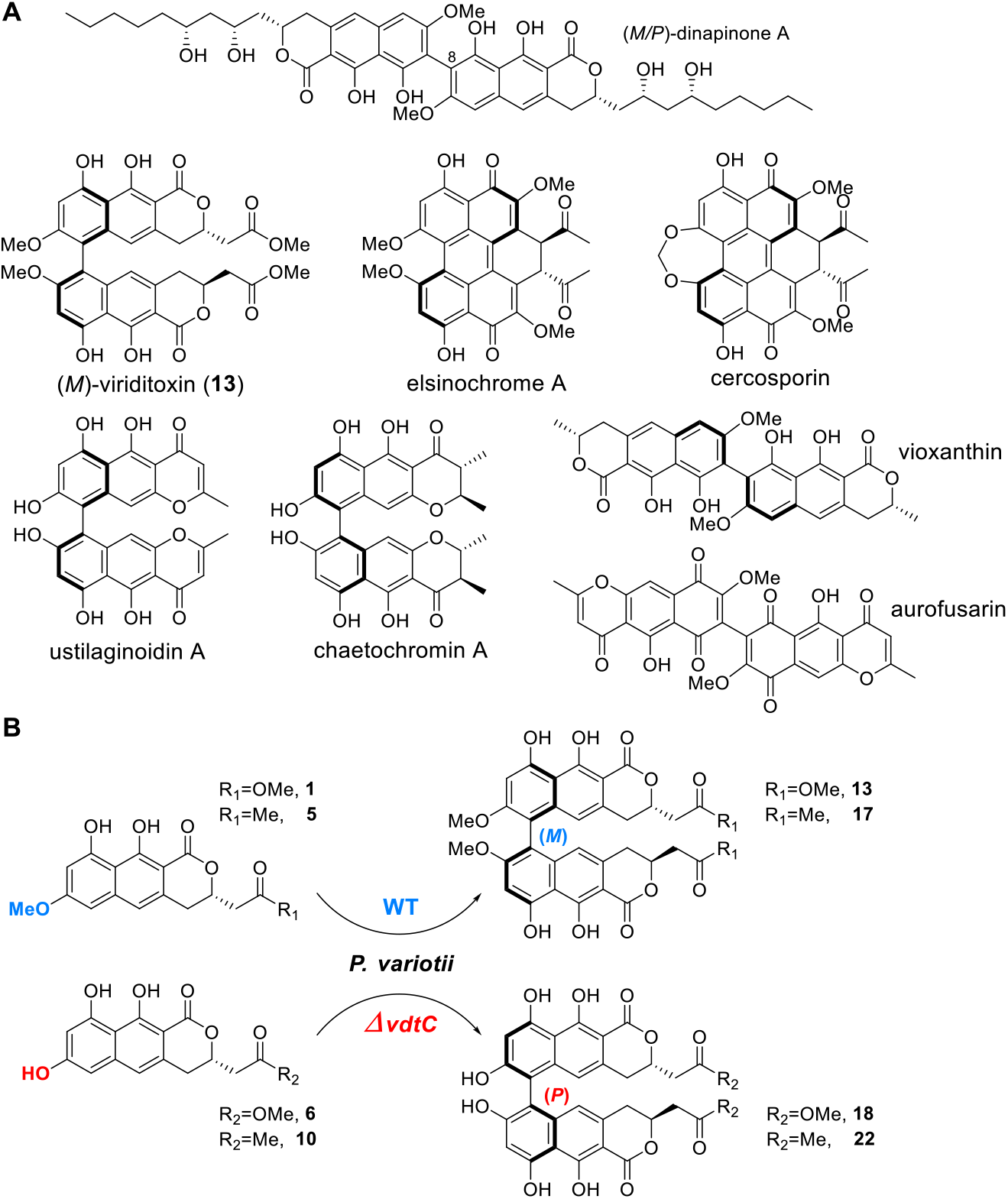
(A) Examples of fungal biaryl products derived from Laccases/MCOs-mediated oxidative coupling. (B) While wild-type *P. variotii* product viriditoxin-related compounds selectively in *M*-form, disruption of *vdtC* resulted in selective accumulation of *P*-form derivatives with loss of *O*-methylation at C7/C7′ position.

We recently characterized an MCO VdtB which is responsible for the final oxidative phenolic coupling step in the biosynthesis of the bisnaphthopyrone (*M*)-viriditoxin in *P. variotii* CBS101075. The heterologous expression-based *in vitro* assays with cell-free lysates showed that with either semi-viriditoxin (**1**), semi-viriditoxin acid (**2**) as substrates, VdtB alone exhibited stringent 6-6′ coupling regioselectivity but yielded coupled products of mixed helical *P/M* chirality at 2:1 ratio. VdtD, a non-catalytic dirigent protein encoded in the cluster, was shown to be involved in controlling the stereoselectivity of VdtB-catalyzed coupling, resulting in the production of (*M*)-viriditoxin in 90% enantiomeric excess (*ee*).^[6c]^ In a parallel study, a targeted gene deletion approach was applied in *P. variotii* to give a comprehensive understanding of viriditoxin biosynthesis.^[6d]^ Curiously, disruption of *vdtC* encoding an *O*-methyltransferase resulted in the production of derivatives with loss of methylation at C7/C7′-OH position as expected, but ECD measurement showed that the derivatives all adopted *P*-chirality, indicating that the substrate structure could potentially have an influence on the stereoselectivity of the VdtB/VdtD-mediated oxidative coupling (Figure 1B). On the other hand, recent characterization of MCOs involved in enantiomeric phenol coupling of *γ*-naphthopyrones highlighted the inherent enantioselectivity of some enzymes in this family, which was previously underestimated.^[6b]^

Inspired by the results above, we set out to test the substrate dependency of the stereoselectivity of VdtB/VdtD system (and the MCO VdtB alone) with various monomeric napthopyrone substrates we obtained during the heterologous reconstruction of the viriditoxin biosynthetic pathway. **1-5, 8**, and **12** were derived from the previous work.^[6c]^ **6, 7**, and **9-11** were further obtained via heterologous expression of different combinations of *vdt* cluster genes in *Aspergillus nidulans*. Variations in these substrates include the presence/absence of *O*-methylation at C7-position, the presence/absence of *Δ^2^* double bond at C3-C4 position, and different moieties at C12 on the side chain (methyl ester/carboxylic acid/ketone). Cell-free lysates of *A. nidulans* individually expressing VdtB and VdtD were used for *in vitro* assays to examine the coupling of the different monomeric substrates (Figure 2).

**Figure 2.**
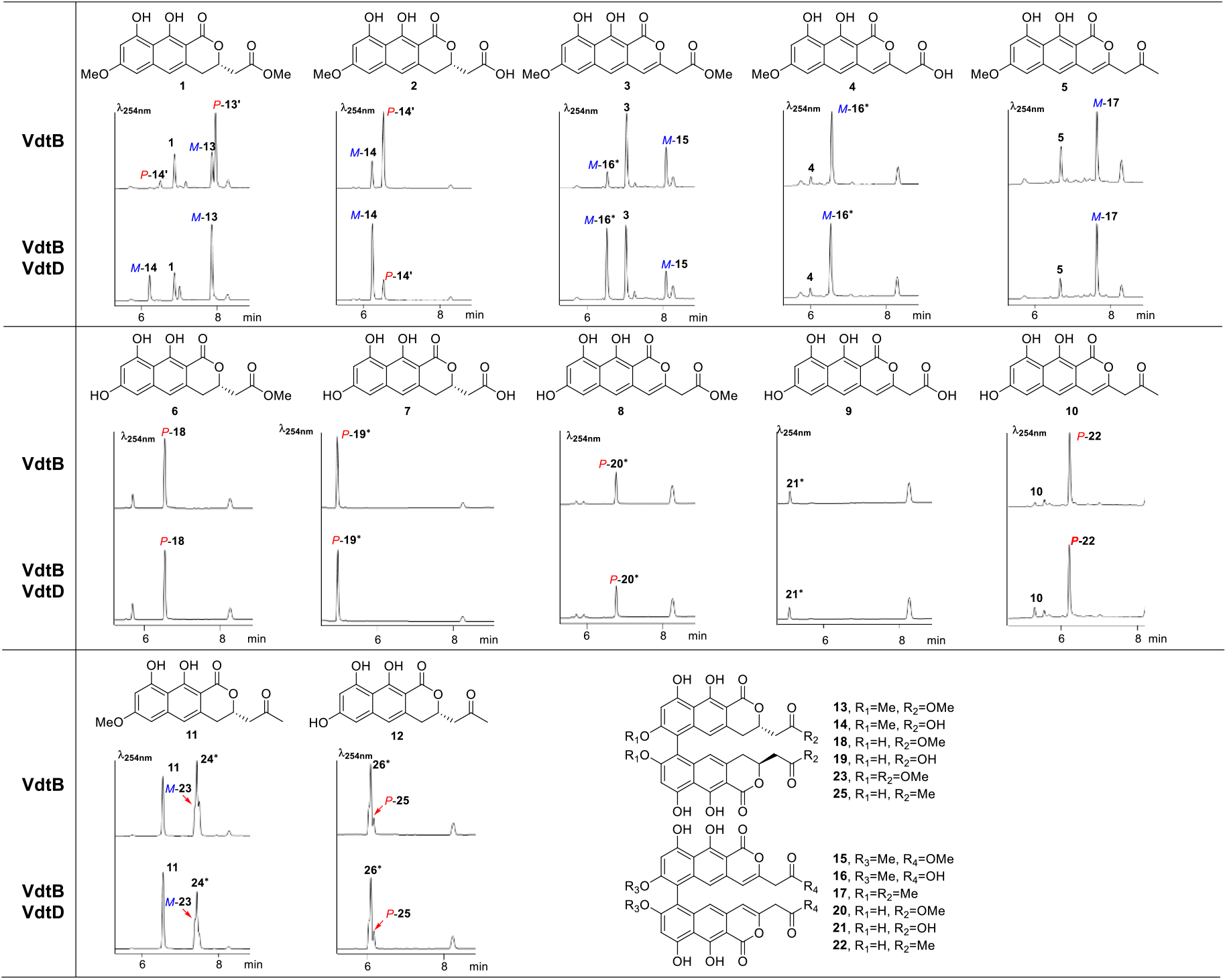
HPLC traces for *in vitro* oxidative coupling of VdtB/VdtD using different substrates. Asterisks indicate compounds of which the structures were not characterized. Compounds labeled with M or P indicated their corresponding helical conformations had been confirmed.

When **1** and **2** were tested, the results remained consistent as previously reported, where the VdtD was shown to direct the stereochemical outcome of the VdtB-catalyzed coupling reaction to produce predominantly bisnaphthopyrone products in *M*-form.^[6c]^ In previous study, *P. variotii* mutants lacking the gene encoding the reductase VdtF produced bisnaphthopyrones *M*-**15** and *M*-**17**, where the pyrone ring is unreduced.^[6d]^ To mimic that, **3, 4** and **5** with the unreduced pyrone ring was subjected to *in vitro* assays using cell-free lysates containing either VdtB and VdtD or VdtB only. When **3** was tested, for both cases (VdtB/D or VdtB only), a peak matching to *M*-**15** could be detected, along with another peak corresponding to the hydrolyzed dimeric derivative **16**, which matched the results when **4** was used as the substrate. Trace amounts of **16** from both sources were acquired for ECD measurement, which confirmed that they were both in the *M*-form (Figure S1). Similar results were observed in the case of **5**, as in both VdtB/D and VdtB only assays, only a single peak matching to dimeric *M***17** was detected as the bisnaphthopyrone product. Chiral HPLC was performed for *M*-**17** standard acquired from *P. variotii ΔvdtF* mutant^[6d]^ and **16** from *in vitro* assay of either **3** or **4** for evaluation of their *ee*. Under the HPLC conditions applied, we could not detect any peak splitting, which likely indicated that the *M*-**17** is enantiomerically pure (Figure S2). The results are surprising, as VdtD is required for the *M*-stereoselectivity of the VdtB-catalyzed oxidative coupling for **1** and **2**, but not for **3** and **4**. The only structural difference between **1** versus **3** and **2** versus **4** is the absence/presence of *Δ^2^* double bond at C3-C4 position on the pyrone ring. The structures of **6** and **7** featured loss of methylation at C7-OH comparing to that of **1** and **2**. The two compounds were applied to the *in vitro* assays to mimic the oxidative coupling in *ΔvdtC* mutant of *P. variotii*.^[6d]^ Consistent with the results of *ΔvdtC* mutants, LC-DAD-MS analysis of the assays with **6** and **7** could only detect a single peak with masses corresponding to their respective dimers (**18** and **19**, respectively), regardless of the presence or absence of VdtD. **18** and **19** were purified likewise from *in vitro* reactions using only VdtB cell lysate, and ECD measurement confirmed that they both adopted a *P*-form (Figure S1). Chiral HPLC was performed for both **18** standard acquired from *P. variotii ΔvdtC* mutantand purified **18** and **19** from *in vitro* assays. No peak separation was detected (Figure S2). The results indicated the lack of methylation at C7-OH on the monomeric substrates altered the stereoselectivity of VdtB to exclusively produce bisnapthopyrone products in the *P*-form. The results also showed that VdtD lost its ability to exert *M*-stereoselectivity on the coupling reactions in the absence of C7-OMe.

To explore the competitive influence between C7-OH and the double bond at C3-C4 position, **8-10** were tested. As shown in Figure 2, single peaks with masses corresponding to the respective dimeric products were detected, regardless of the presence/absence of VdtD. The standards of **20** and **21** were not available from previous study. However, given the well-acknowledged 6,6′ regioselectivity of VdtB and its homologs,^[6c, 6e]^ they were also suggested to be 6,6′-coupling dimers. **22** matched to the *P*-**22** standard acquired from the heterologous expression of *vdtAB* in *A. nidulans*. Only **20** was successfully purified from scaled-up *in vitro* reaction and its chiral HPLC was performed together with *P*-**22** standard. While no peak splitting was detected in trace of **20**, one tiny peak corresponding to potentially *M*-**22** was observed, resulting in an *ee* of 85% in *P*-**22** standard (Figure S2). The results suggested a stronger influence of C7-OH over that of C3-C4 double bond on the stereoselectivity of VdtB, and that VdtD has no effect on the stereoselectivity of VdtB for these substrates.

VdtE is a Baeyer-Villiger monooxygenase responsible for the generation of methyl ester from the ketone side-chain of the monomeric intermediates **11**.^[6c]^ The gene deletion study showed that Δ*vdtE* mutants produced exclusively bisnaphthopyrone *M*-**23**, which corresponds to the dimer of **11**. Interestingly, when **11** was used as a substrate for VdtB-only cell-free lysate assay, it resulted in three peaks in LC-DAD-MS trace, which all have identical UV-vis spectrum and mass. Two of the peaks matched to the corresponding standards for atropisomers *M*/*P*-**23** (*M*-**23** was obtained from WT *P. variotii* while *P*-**23** was obtained from heating of *M*-**23**), but the third peak **24** did not match to any of the dimeric standards we have. Similar results were observed when **12** was tested for VdtB-only assay, where besides *M*/*P*-**25**, an additional peak **26** with identical UV-vis spectrum and mass was detected. Although further structural characterizations were needed for **24** and **26**, it seems that the only reasonable explanation is that they are bisnaphthopyrone products derived from a non-6,6′-coupling reaction. This raised the question about the timing of VdtF-catalyzed C3-C4 reduction in the biosynthetic pathway of viriditoxin, as the observation that Δ*vdtE* mutant produced exclusively *M*-**23** might be a result of coupling of the monomeric substrate **5** to *M*-**17** followed by reduction of the dimer product to *M*-**23**. The results also suggest that incorrect substrate could lead to the loss of the regio-selective control of VdtB. In terms of stereoselectivity, assays using cell-free lysate of VdtB-only produced the *M/P*-**23** in approximately 1:1 ratio, while the addition of VdtD resulted in the production of higher level of the *M*-form of **23**. For substrate **12**, VdtB-only assay resulted in higher level of the *M*-form of **25**, and the addition of VdtD resulted in no change in the *M/P* ratio.

Our results clearly showed that the stereoselectivity of VdtB could be strongly influenced by the structural modifications on the substrates. With the modification of certain moieties in the substrate structure, the stereoselective control of VdtB could be reversed (as shown by **1-2** *vs.* **3-5** *vs.* **6-7** *vs.* **8-10**). The presence of C3-C4 double bond changed the stereoselectivity of VdtB to the overall *M*-form, while the loss of methylation at C7-OH resulted in a *P*-preferred coupling. Such substrate effects could also be observed in the ability of VdtD to exert stereoselective control of the VdtB-catalyzed coupling. *In vitro* assays clearly showed that the ability of VdtD to mediate the *M*-stereoselectivity on VdtB is strong in the coupling of **1** but completely lost in the coupling of **6-10**, probably due to the lower binding affinity to these substrates. For **3-5**, it is difficult to determine whether VdtD was functional as the coupling resulted in products in the *M*-form regardless of the absence or presence of VdtD. In the case of **2**, however, we observed that the efficiency of such control by VdtD dropped as we observed a slightly lower ratio of *M*/*P*-**14** compared to *M*/*P*-**13** when **1** is used as the substrate. To examine this, we tested if the concentration of VdtD could have an effect on the ratio of the atropisomer products. We found that in order to obtain comparable *M*-selective coupling for **2** when compared with that of **1**, a higher concentration of VdtD was required (Figure 3). This substrate-dependent nature of the stereoselectivity we observed in the binary VdtB/VdtD-mediated oxidative coupling led us to propose a VdtB-ligand-VdtD complex model whereby the stereoselective outcome of the coupling reaction is a result of the interplay of interactions between the substrate, and both VdtB and VdtD, which have different substrate affinities.

**Figure 3.**
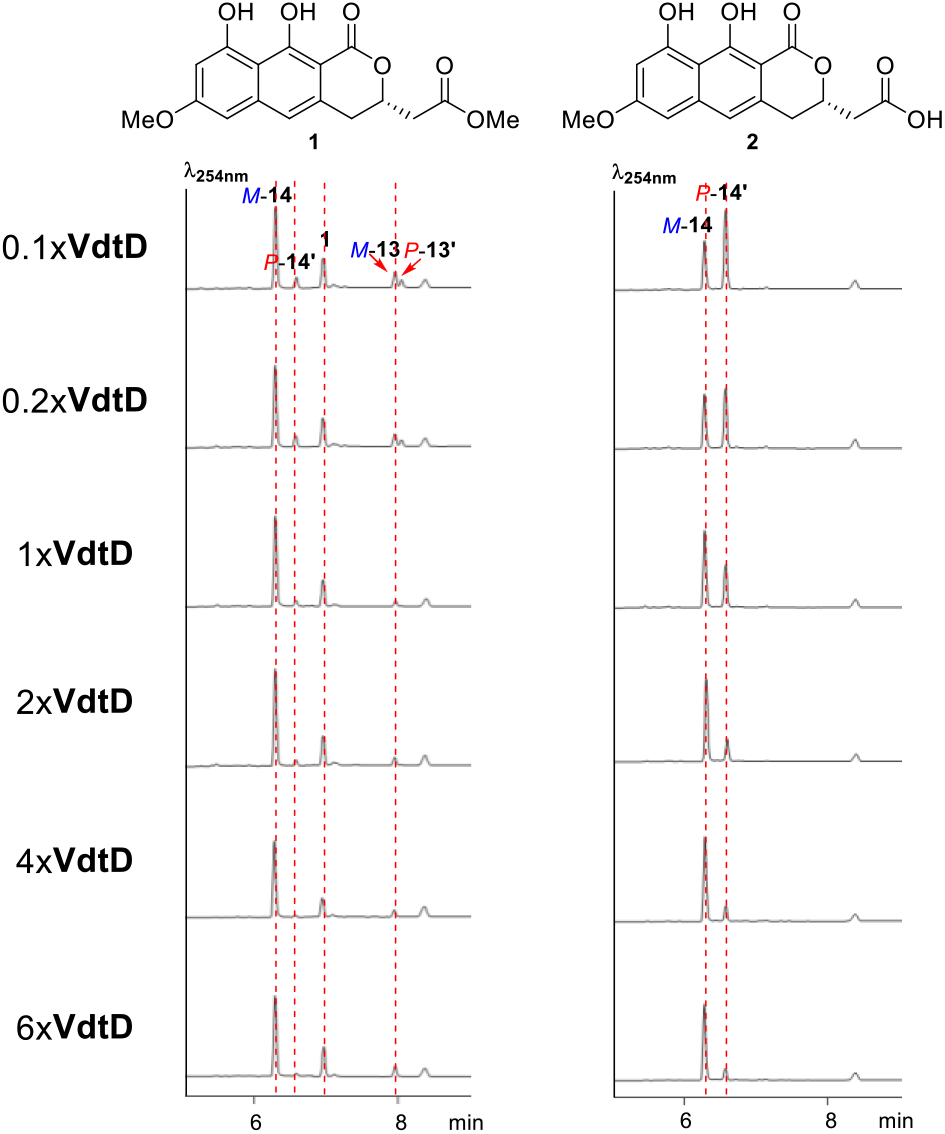
The stereoselective control of VdtD over VdtB on coupling of **1** and **2**. The endogenous hydrolase activity increased when larger amounts of mycelium were used for preparing cell-free lysates. This endogenous hydrolase specifically acts on *M*-13 (Figure S6). The difference between the ratio of *M*-14:*P*-14 suggested that hydrolysis proceeded after the oxidative coupling catalyzed by VdtB.

All the recently characterized LMCOs from fungal biaryls biosynthetic gene clusters (BGCs) shared significant protein sequence similarities. *Tp*MCO from *Talaromyces pinophilus* was characterized to catalyze the 8,8′-regioselective coupling in the biosynthesis of dinapinones (Figure 1).^[6a]^ Four laccases, namely UstL, CheL, AshL, and MytL, were recently characterized to be involved in the stereoselective coupling of γ-naphthopyrones.^[6b]^ Using nor-rubrofusarin as the substrate, CheL, AshL, and MytL showed stringent *P*-selective coupling ability. UstL, on the other hand, showed flexible stereoselectivity. VdtB shares 53% protein sequence identity with *Tp*MCO and 52-55% identity with the four MCOs mentioned above. Their catalytic features also have some resemblances. Under most *in vitro* conditions and substrate derivatives tested, the LMCOs exhibited consistent regioselectivity, but the stereoselectivity of the LMCOs could vary depends on the conditions or substrates tested. For example, dinapinones A1 and A2 were initially isolated as atropisomers with an *M*:*P* ratio of close to 1:1 *in vivo*, while purified *Tp*MCE showed a *P*-preferred coupling with an *M*:*P* ratio of 1:2.5 *in vitro*.^[6a, 7]^ When a non-native substrate semi-vioxanthin was used for assaying Av-VirL, a homolog of VdtB from *Aspergillus viridinutans*, the observed stereochemical outcome of the coupling reaction favoured the *P*-form instead of the expected *M*-form observed for viriditoxin.^[6e]^ Remarkable flexibility was also observed for the stereoselectivity of UstL depends on its concentration, where only biaryls in the *M*-form could be isolated from its original host *Ustilaginoidea virens*.^[6b, 8]^ The results in this work showed that variations in substrate structures could have a major impact on the stereoselectivity of VdtB, and we hypothesize that this property is potentially applicable to other LMCOs.

Thus, this study together with the recent work clearly demonstrated that fungal LMCOs have the potential to catalyze oxidative coupling in a stereoselective manner.^[6b]^ However, the presence of accessory protein, such as the dirigent protein VdtD, could affect the stereoselectivity of the LMCO-catalyzed coupling. VdtD was initially predicted to be non-catalytic *α/β*-hydrolase because of a key mutation of S228D in the classic catalytic triad of serine hydrolases. A mutant of VdtD carrying D228S mutation was also generated here to restore the serine in the catalytic triad. Interestingly, the mutant showed same level of stereoselective influence over VdtB and no observable hydrolase activity for the methyl ester moiety in the monomeric substrates, suggesting that the whole protein has undergone neofunctionalization (Figure S6). There were suggestions that there is at least another family of accessory protein that accompanies fungal LMCOs, which is known as fasciclin domain-containing proteins (FACs). The presence of such non-catalytic FACs has been identified in multiple LMCOs-mediated biaryl BGCs, and their importance in the biosynthesis has been highlighted (Figure S5).^[6e, 9]^ In a previous study about the biosynthesis of ustilaginoidins, a FAC was also present in the BGC identified, but its function was uncharacterized.^[10]^ We hypothesize that their role might resemble that of VdtD in viriditoxin biosynthesis.

Another interesting hypothesis from this study is that it suggests, for the exclusive production of (*M*)-viriditoxin to occur in *P. variotii*, a stringent order of the biosynthetic steps shall be followed in the pathway. This is because the *in vitro* assay results suggested that VdtB possess considerable substrate flexibility (Figure S7-S8). For example, VdtC-catalyzed C7-OH methylation has to occur prior to VdtB-catalyzed oxidative coupling for the formation of the *M*-bisnapthopyrones.. In the previous study, *in vitro* assay of VdtE also suggested that in order to remain its regioselectivity, **5** but not **11** is the correct substrate.^[6c]^ This makes us wonder if there exists a multi-protein complex (at least VdtC, VdtB, and VdtD in the case of *vdt* cluster), although the compartmentalization of enzymes in different compartments in fungal SMs biosynthesis could not be excluded.^[11]^ In summary, this work highlights the substrate-dependent stereoselectivity of VdtB and identifies the key moieties that could influence the stereoselectivity of VdtB. It was demonstrated that VdtB could accept a range of structurally related naphthopyrone substrates. For certain substrates, the coupling reactions catalyzed by VdtB were highly stereoselective, thus making VdtB a potentially powerful enzymatic tool for stereoselective phenol oxidative coupling. The dirigent protein VdtD provides another layer of stereoselective control for VdtB, although the substrate scope for VdtD is more limited. Despite the widespread use of fungal MCOs in industries and their biotechnological potentials, the detailed mechanisms of MCOs-mediated oxidative coupling reactions are currently under-studied. Future exploration of the molecular basis of VdtB/VdtD-mediated regio- and stereoselective oxidative coupling could provide more insights into the application of these powerful enzymes.

## Experimental Section

Experimental details, supplementary figures, and spectroscopic data could be found in supplementary information which could be found at https://doi.org/10.1101/846196

## Acknowledgements

Y.-H.C. and G.R.F. are Australian Research Council (ARC) Future Fellows. This study was supported by an Australian Research Council (ARC) Discovery Project (DP170100228). NMR and HR-MS analyses were performed at the UWA Centre for Microscopy, Characterisation and Analysis (CMCA). JH is recipient of Australian International Postgraduate Research Scholarship.

