## supporting information for "The substrate-dependent stereoselectivity of the multicopper oxidase (MCO)-catalyzed oxidative coupling in the biosynthesis of the bisnaphthopyrone viriditoxin"

### Experimental procedures section

#### Strains and culture conditions

Glucose minimal medium (GMM, 10 g Glucose, 6g $NaNO_3$, 1.52 g $K_2HPO_4$, 0.52 g KCl, 0.52 g $MgSO_4·7H_2O$, 22 mg $ZnSO_4·7H_2O$, 11 mg $H_3BO_3$, 5 mg $MnCl_2·4H2O$, 1.6 mg $FeSO_4·7H2O$, 1.6 mg $CoCl_2·5H_2O$, 1.6 mg $CuSO_4·5H_2O$, 1.1 mg $(NH_4)_6Mo_7O_{24}·4H_2O$, 50 mg $Na_4$EDTA in 1L, adjusted to pH 6.5) was used for culturing of *A. nidulans*. Uracil/Uridine, riboflavin, and pyridoxine with suggested concentration are supplemented if necessary. In terms of *A. nidulans* protoplast transformation, Stabilized minimal medium (SMM, GMM supplemented with 1.2 M sorbitol) was used.

For small-scale metabolic profile analysis of *A. nidulans* culture, spores with a concentration of ~$10^8$/L were inoculated into 50ml GMM media. Cultures were grown at 37 °C/180rpm for 18h. 2.5ml/L of cyclopentanone was added to induce the expression of $P_{alc}$ promoters.[1] Cultures were incubated at 25 °C/180rpm for another 4 days for the accumulation of secondary metabolites. For each combination, three separate transformants were peaked as replicates. For isolation of target compounds, scaled-up cultures were performed under the same conditions.

#### Construction of Yeast-Fungal Artificial Chromosomes (pYFACs) for heterologous expression in *A. nidulans*

*A. nidulans* heterologous expression vectors (pYFAC-CH1/2/4/5) were constructed in previous work. Yeast homologous recombination was used for the construction of YFACs in this work. Full-length genomic fragments of *vdt1-6* were amplified with primers designed and cloned into corresponding vectors (Table S1) as previously described.[2] Yeast miniprep of corrected transformants was performed with Zymoprep™ Yeast Plasmid Miniprep I Kit. 1 ul from the miniprep was used for *E. coli* NEB10-beta® electroporation. Samples with correct digestion pattern were used for *A. nidulans* transformation as previously described.[1]

#### Sample preparation and Metabolic profile analysis

For *A. nidulans* cultures, extracts from both mycelium and media were analyzed. Mycelium was collected by filtration and extracted with acetone. Media was extracted with either ethyl acetate/methanol/acetic acid (89:10:1). Crude extracts were dried down and redissolved in methanol for LC-DAD-MS analysis. For *P. variotii* cultures, agar culture was chopped into small pieces and extracted with methanol under sonication condition. Crude extracts were filtered, dried and redissolved in methanol for LC-DAD-MS analysis.

LC-DAD-MS analysis was performed with Agilent 1260 liquid chromatography (LC) system coupled to a diode array detector (DAD) and an Agilent 6130 quadrupole mass spectrum (MS) with an ESI source. For analytical purpose, a Kinetex C18 column (2.6 µm, 2.1 mm i.d. x 100 mm; Phenomenex) was used. The mobile phase gradient of eluent B (acetonitrile with 0.1% formic acid) started at 5% and gradually increased to 95% over 10 mins at a flow rate of 0.75 ml/min.

Chiral HPLC analysis was performed with Chiralcel OJ-R cellulose column (5 μm, 4.6 mm i.d. x 150 mm, Dicel). The mobile phase of eluent B (acetonitrile with 0.1% formic acid) was run with an isocratic method varying from 30% to 60% with a flow rate of 0.5 ml/min.

**Compound isolation and structural characterization.**

Ethyl acetate/methanol/acetic acid (89:10:1) extraction method was used for isolation of **4**-**13** from *A. nidulans* culture medium. Ethyl acetate/methanol (90:10) was used for extraction from *P. variotii* CYA culture. Crude extracts were dried *in vacuo* and then fractionated on a Reveleris flash chromatography system (Grace) using a dichloromethane /methanol gradient on a Reveleris HP silica flash cartridge. Fractions containing the target compound were combined for further purification using a semi-prep HPLC with a C18 column (Agilent, 5 μm, 21.2 × 150 mm).

For structural characterization, nuclear magnetic resonance (NMR) spectra were collected on Bruker Avance IIIHD 500MHz/600MHz NMR spectrometer. Chloroform-*d*, DMSO-$d_6$, acetonitrile-$d_3$, and acetone-$d_6$ were used as solvents. Electronic circular dichroism (ECD) spectra were recorded on a JASCO J-810 spectropolarimeter, with acetonitrile as solvent. The axial chirality of dimeric compounds isolated was determined by comparing with both the ECD spectrum of (*M*)-viriditoxin standard as previously described.[2]

Standards of **1**-**5**, **10**, **13**-**15**, **17**-**18**, and **23**-**24** were derived from previous works. [2, 3] Compounds **6**-**9**, **11**-**12** and **22** were isolated in this work.

**Preparation of cell-free extracts for VdtB and VdtD**

pYFAC-10/11 expressing *vdtB*/*vdtD* was transformed into *A. nidulans*. Mycelium was harvested after 3-days induction, frozen with liquid nitrogen and grinded into a fine powder with pestle and mortar. The powder was resuspended in 50 mM citrate buffer pH 5.0 supplemented with 2 mM DTT and lysed by sonication on ice. Cellular debris was removed by centrifugation at 17000 rpm, 4 °C for 40 min and the supernatant was used directly for *in vitro* assays.

***in vitro*** **enzymatic reaction assay for VdtB, VdtD**

All *in vitro* reactions were performed at 100 μl scale in 50 mM citrate buffer, pH 5.0.[2] 40 μl VdtB cell-free lysate/purified fraction and an equal volume of VdtD cell-free lysate/purified fraction was added together with 2 mM substrates**.** 0.1% Triton X-100 was also added to increase compound solubility. The reaction was carried out at 30 °C, 400 rpm and incubated for 5 hours. The reactions were quenched with 200 μl of ethyl acetate/methanol/acetic acid (89:10:1). The organic phase was separated, dried, redissolved in 50 μl of methanol, and analyzed with LC-DAD-MS as previously described.

### Supplementary Tables

#### Table S1. List of constructs used in this work.

| Construct name | Description | Source |
| --- | --- | --- |
| pYFAC-CH1 | *AMA1*; *CEN/ARS*; *ColE1 ori*; *pyrG*; *URA3*; $Amp^{R}$; $P_{alcA}$. | |
| pYFAC-CH4 | *AMA1*; *CEN/ARS*; *ColE1 ori*; *pyrO*; *URA3*; $Amp^{R}$; $P_{alcA}$-T1-$P_{alcSM}$-T2-$P_{aldA}$. | |
| pYFAC-CH5 | *AMA1*; *CEN/ARS*; *ColE1 ori*; *ribo*; *URA3*; $Amp^{R}$; $P_{alcA}$-T1-$P_{alcSM}$. | |
| pYFAC-1 | pYFAC-CH1 expressing *vdtA* | [2] |
| pYFAC-2 | pYFAC-CH5 expressing *vdtB* | [2] |
| pYFAC-3 | pYFAC-CH4 expressing *vdtC* | [2] |
| pYFAC-4 | pYFAC-CH4 expressing *vdtC* and *vdtD* | [2] |
| pYFAC-5 | pYFAC-CH4 expressing *vdtC* and *vdtE* | [2] |
| pYFAC-6 | pYFAC-CH4 expressing *vdtC* and *vdtF* | [2] |
| pYFAC-7 | pYFAC-CH4 expressing *vdtC*, *vdtD* and *vdtE* | [2] |
| pYFAC-8 | pYFAC-CH4 expressing *vdtC-F* | [2] |
| pYFAC-10 | pYFAC-CH1 expressing *vdtD* fused with 6xHis-tag | [2] |
| pYFAC-11 | pYFAC-CH1 expressing *vdtB* fused with MBP-tag | This work |
| pYFAC-12 | pYFAC-CH4 expressing *vdtE* | This work |
| pYFAC-13 | pYFAC-CH4 expressing *vdtF* | This work |
| pYFAC-14 | pYFAC-CH4 expressing *vdtC* and *vdtF* | This work |
| pYFAC-15 | pYFAC-CH5 expressing *vdtB* and *vdtD* | This work |

#### Table S2. Structural information of 6 (DMSO-$d_6$).

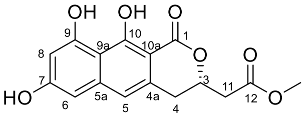

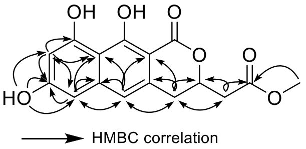

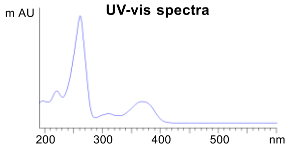

| Carbon No. | $^{13}$C NMR | $^{1}$H NMR (ppm, multi, *J*) | gCOSY | HMBC |
|---|---|---|---|---|
| 1 | .- | - | - | - |
| 2 | - | - | - | - |
| 3 | 75.7 | 4.95 (1H, m) | - | - |
| 4 | 31.7 | 3.05 (1H, dd, *J*=2.90, 13.40)<br>2.99 (1H, dd, *J*=3.01, 13.15) | - | 3, 4a, 5, 10a, 11 |
| 4a | 133.1 | - | | - |
| 5 | 115.1 | 6.87 (1H, s) | - | 4, 5a, 6, 9a, 10a |
| 5a | 140.7 | - | - | - |
| 6 | 101.9 | 6.53 (1H, d, 1.70) | - | 5, 5a, 7, 8, 9a |
| 7 | 160.9 | - | - | - |
| 7-OH | - | 10.22 (1H, s) | - | 6, 7, 8 |
| 8 | 101.8 | 6.39 (1H, d, 1.75) | - | 6, 7, 9, 9a |
| 9 | 158.3 | - | - | - |
| 9a | 107.1 | - | - | - |
| 10 | 162.6 | - | - | - |
| 10a | 99.1 | - | - | - |
| 11 | 38.8 | 2.92 (1H, dd, *J*=4.00, 13.55)<br>2.84 (1H, dd, *J*=6.75, 13.55) | - | 3, 4, 12 |
| 12 | 170.0 | - | - | - |
| 12-OMe | 51.7 | 3.66 (3H, s) | | 12 |

#### Table S3. Structural information of 7 (MeOD-$d_4$).

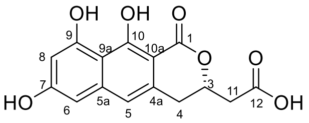

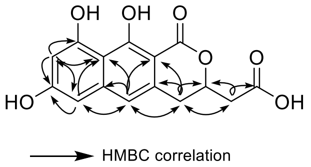

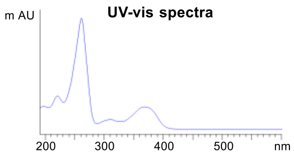

| bon No. | $^{13}$C NMR | $^{1}$H NMR (ppm, multi, *J*) | gCOSY | HMBC |
|---|---|---|---|---|
| 1 | 172,3 | - | - | - |
| 2 | - | - | - | - |
| 3 | 77.8 | 4.99 (1H, m) | - | - |
| 4 | 33.3 | 3.00 (1H, dd, *J*=2.40, 13.35)<br>3.10 (1H, dd, *J*=9.30, 13.05) | - | 3, 4a, 5, 10a, 11 |
| 4a | 134.2 | - | | - |
| 5 | 116.9 | 6.84 (1H, s) | - | 4, 5a, 6, 9a, 10a |
| 5a | 142.4 | - | - | - |
| 6 | 103.2 | 6.51 (1H, d, 1.75) | - | 5, 7, 8, 9a |
| 7 | 162.6 | - | - | - |
| 8 | 102.9 | 6.37 (1H, d, 1.80) | - | 6, 7, 9, 9a |
| 9 | 159.9 | - | - | - |
| 9a | 108.3 | - | - | - |
| 10 | 163.7 | - | - | - |
| 10a | 99.6 | - | - | - |
| 11 | 40.3 | 2.80 (1H, dd, *J*=4.65, 13.60)<br>2.84 (1H, dd, *J*=6.15, 13.55) | - | 3, 4, 12 |
| 12 | 173.2 | - | - | - |

#### Table S4. Structural information of 8 (DMSO-$d_4$).

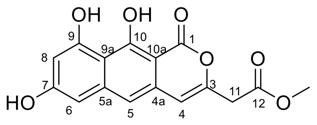

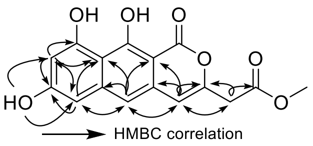

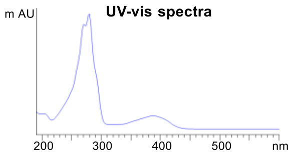

| Carbon No. | $^{13}$C NMR | $^{1}$H NMR (ppm, multi, *J*) | gCOSY | HMBC |
|---|---|---|---|---|
| 1 | .166.1 | - | - | - |
| 2 | - | - | - | - |
| 3 | 148.2 | - | - | - |
| 4 | 106.6 | 6.59 (1H, s) | 11 | 3, 4a, 5, 10a, 11 |
| 4a | 131.0 | - | | - |
| 5 | 112.1 | 7.02 (1H, s) | - | 4, 5a, 6, 9a, 10a |
| 5a | 141.6 | - | - | - |
| 6 | 101.6 | 6.60 (1H, d, 2.15) | - | 5, 7, 8, 9a |
| 7 | 161.0 | - | - | - |
| 7-OH | - | 10.32 (1H, s) | - | 6, 7, 8 |
| 8 | 101.9 | 6.43 (1H, d, 2.05) | - | 6, 7, 9, 9a |
| 9 | 158.5 | - | - | - |
| 9a | 107.4 | - | - | - |
| 10 | 162.8 | - | - | - |
| 10a | 96.7 | - | - | - |
| 11 | 38.1 | 3.70 (2H, s) | 4 | 3, 4, 12 |
| 12 | 169.0 | - | - | - |
| 12-OMe | 52.2 | 3.68 (3H, s) | | 12 |

#### Table S5. Structural information of 9 (DMSO-$d_4$).

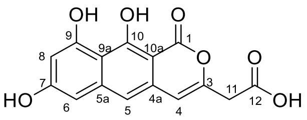

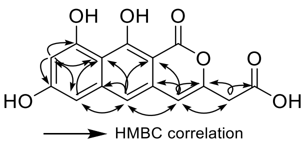

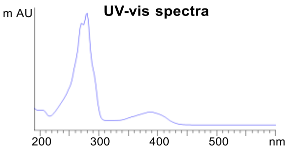

| Carbon No. | $^{13}$C NMR | $^{1}$H NMR (ppm, multi, *J*) | gCOSY | HMBC |
|---|---|---|---|---|
| 1 | .166.2 | - | - | - |
| 2 | - | - | - | - |
| 3 | 149.2 | - | - | - |
| 4 | 106.6 | 6.56 (1H, s) | - | 3, 4a, 5, 10a, 11 |
| 4a | 131.2 | - | | - |
| 5 | 111.8 | 7.02 (1H, s) | - | 4, 5a, 6, 9a, 10a |
| 5a | 141.6 | - | - | - |
| 6 | 101.5 | 6.59 (1H, d, 2.10) | - | 5, 7, 8, 9a |
| 7 | 162.0 | - | - | - |
| 7-OH | - | 10.29 (1H, s) | - | 6, 7, 8 |
| 8 | 101.8 | 6.42 (1H, d, 2.10) | - | 6, 7, 9, 9a |
| 9 | 158.6 | - | - | - |
| 9a | 107.4 | - | - | - |
| 10 | 163.0 | - | - | - |
| 10a | 96.7 | - | - | - |
| 11 | 38.5 | 3.57 (2H, s) | - | 3, 4, 12 |
| 12 | 170.0 | - | - | - |

#### Table S6. Structural information of 12 (DMSO-$d_4$).

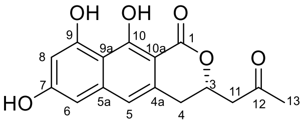

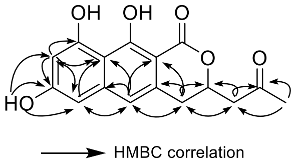

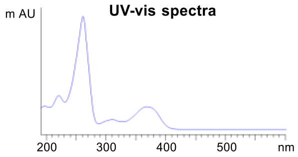

| Carbon No. | $^{13}$C NMR | $^{1}$H NMR (ppm, multi, *J*) | gCOSY | HMBC |
|---|---|---|---|---|
| 1 | 171.3 | - | - | - |
| 2 | - | - | - | - |
| 3 | 75.6 | 5.08 (1H, m) | - | - |
| 4 | 31.9 | 2.9 (1H, dd, -)[a]<br>3.1 (1H, dd, -)[a] | - | 3, 4a, 5, 10a, 11 |
| 4a | 133.3 | - | | - |
| 5 | 115.0 | 6.89 (1H, s) | - | 4, 5a, 6, 9a, 10, 10a |
| 5a | 140.6 | - | - | - |
| 6 | 101.8 | 6.53 (1H, d, 2.58) | - | 5, 8, 9a |
| 7 | 163.2 | - | - | - |
| 7-OH | - | 10.21 (1H, s) | - | 6, 7, 8 |
| 8 | 101.7 | 6.38 (1H, d, 2.64) | - | 6, 7, 9, 9a |
| 9 | 158.2 | - | - | - |
| 9a | 107.0 | - | - | - |
| 10 | 162.5 | - | - | - |
| 10a | 98.6 | - | - | - |
| 11 | 47.3 | 2.9 (1H, dd, -)[a]<br>3.1 (1H, dd, -)[a] | | 3, 4, 12 |
| 12 | 205.2 | - | - | - |
| 13 | 30.2 | 2.17 (3H, s) | - | 11, 12 |

[a] unable to calculate *J* due to peak overlapping

#### Table S7. Structural information of 22 (acetone-$d_3$).

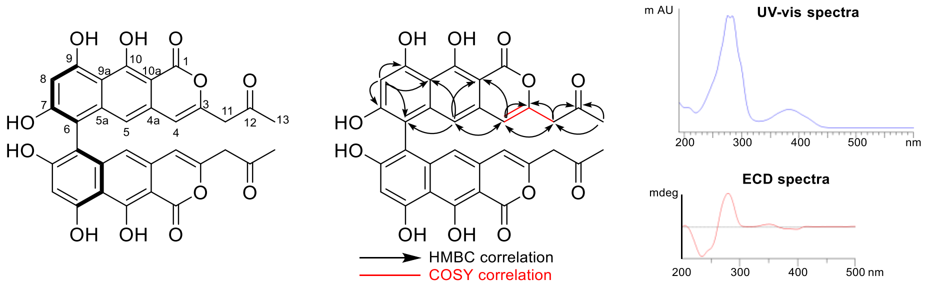

| Carbon No. | $^{13}$C NMR | $^{1}$H NMR (ppm, multi, *J*) | gCOSY | HMBC |
|---|---|---|---|---|
| 1 | .168.3[a] | - | - | - |
| 2 | - | - | - | - |
| 3 | 150.7 | - | - | - |
| 4 | 108.4 | 6.34 (1H, s) | 11 | 3, 5, 10a, 11 |
| 4a | 132.7[a] | - | | - |
| 5 | 111.5[b] | 6.50 (1H, s) | - | 4, 6, 9a, 10a |
| 5a | 142.4[a] | - | - | - |
| 6 | 109.2[b] | - | - | - |
| 7 | 160.2[b] | - | - | - |
| 7-OH | - | - | - | - |
| 8 | 102.8 | 6.63 (1H, d, 2.04) | - | 6, 7, 9, 9a |
| 9 | 161.1 | - | - | - |
| 9-OH | - | - | - | - |
| 9a | 106.8[b] | - | - | - |
| 10 | - | - | - | - |
| 10-OH | - | - | - | - |
| 10a | 98.1 | - | - | - |
| 11 | 47.8 | 3.63 (2H, s) | 4 | 3, 4, 12 |
| 12 | 202.8 | - | - | - |
| 13 | 29.6 | 2.19 (3H, s) | - | 11, 12 |

[a] $^{13}$C signals have no HMBC correlations. The chemical shifts were assigned based on data of other compounds.

[b] Weak signals in $^{13}$C spectrum.

### Supplementary Figures

#### Figure S1. ECD spectrum of *M*-16, *P*-18, *P*-19, and *P*-20

i) *M*-**16** purified from VdtB catalyzed *in vitro* coupling of **3**; ii) *M*-**16** purified from VdtB catalyzed *in vitro* coupling of **4**; iii) *P*-**18** purified from VdtB catalyzed *in vitro* coupling of **6**; iv) *P*-**19** purified from VdtB catalyzed *in vitro* coupling of **7**; v) *P*-**20** purified from VdtB catalyzed *in vitro* coupling of **8**.

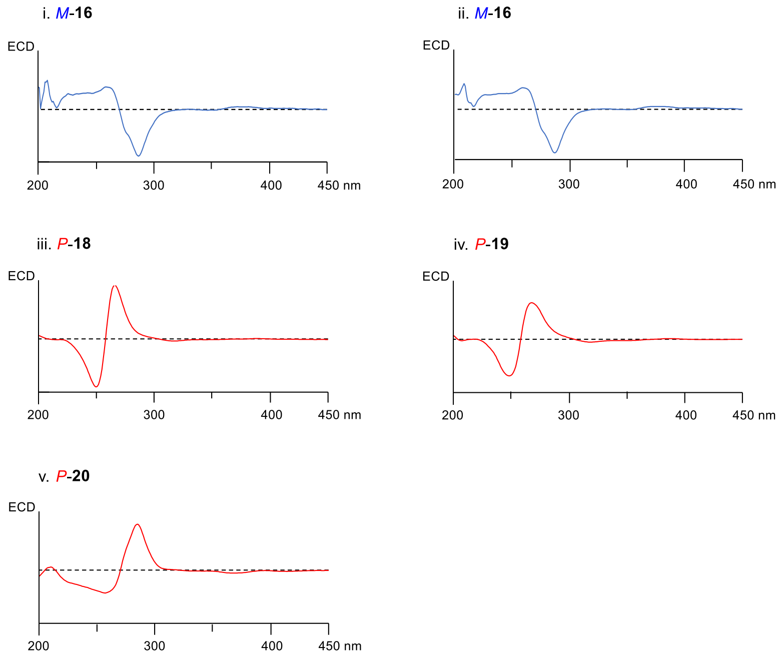

**Figure S2. Chiral HPLC traces for *M*-15, *M*-16, *M*-17, *P*-18, *P*-19, *P*-21, and *P*-22**

i) *M*-**15** isolated from *P. variotii* △*vdtF* mutant;

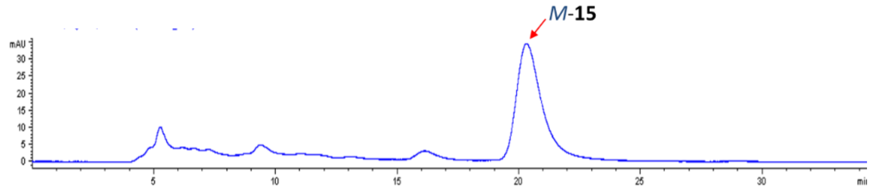

ii) *M*-**16** purified from VdtB catalyzed *in vitro* coupling of **3**;

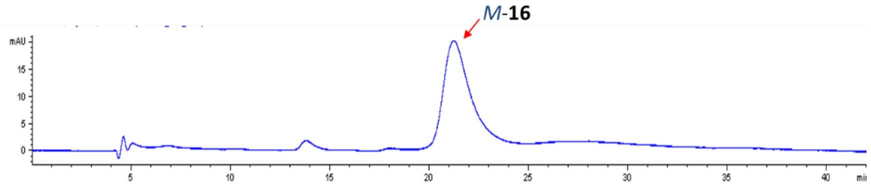

iii) *M*-**17** isolated from *P. variotii* △*vdtF* mutant;

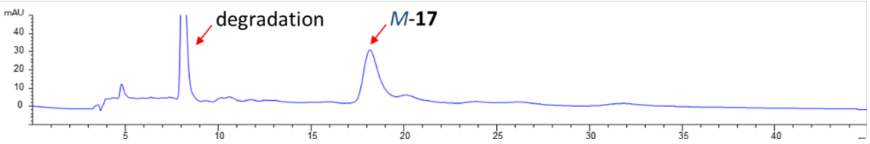

iv) *P*-**18** purified from both VdtB catalyzed *in vitro* coupling of **6** and *P. variotii* △*vdtC* mutant;

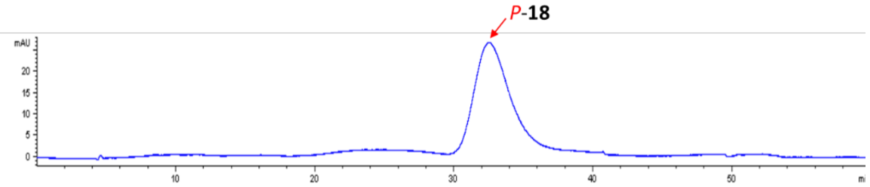

v) *P*-**19** purified from VdtB catalyzed *in vitro* coupling of **7**;

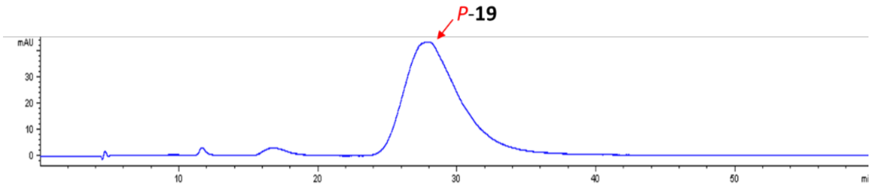

vi) *P*-**20** purified from VdtB catalyzed *in vitro* coupling of **8**;

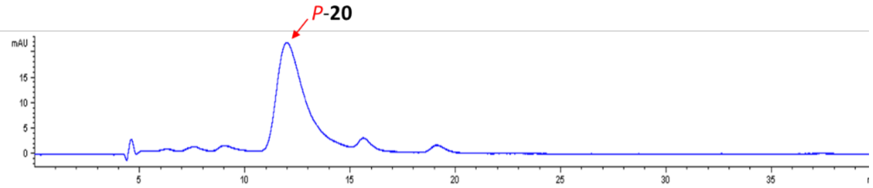

vii) *P*-**22** was isolated from heterologous expression of *vdtAB* in *A. nidulans*.

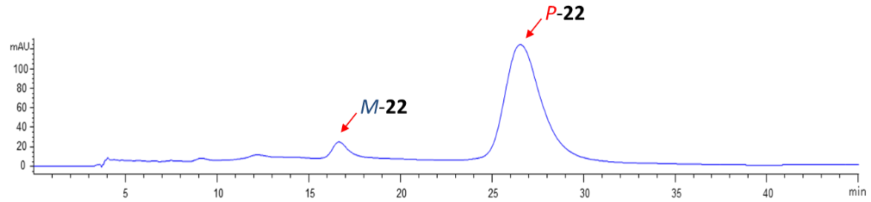

### Figure S3. *In vitro* assay results with standards or EIC.

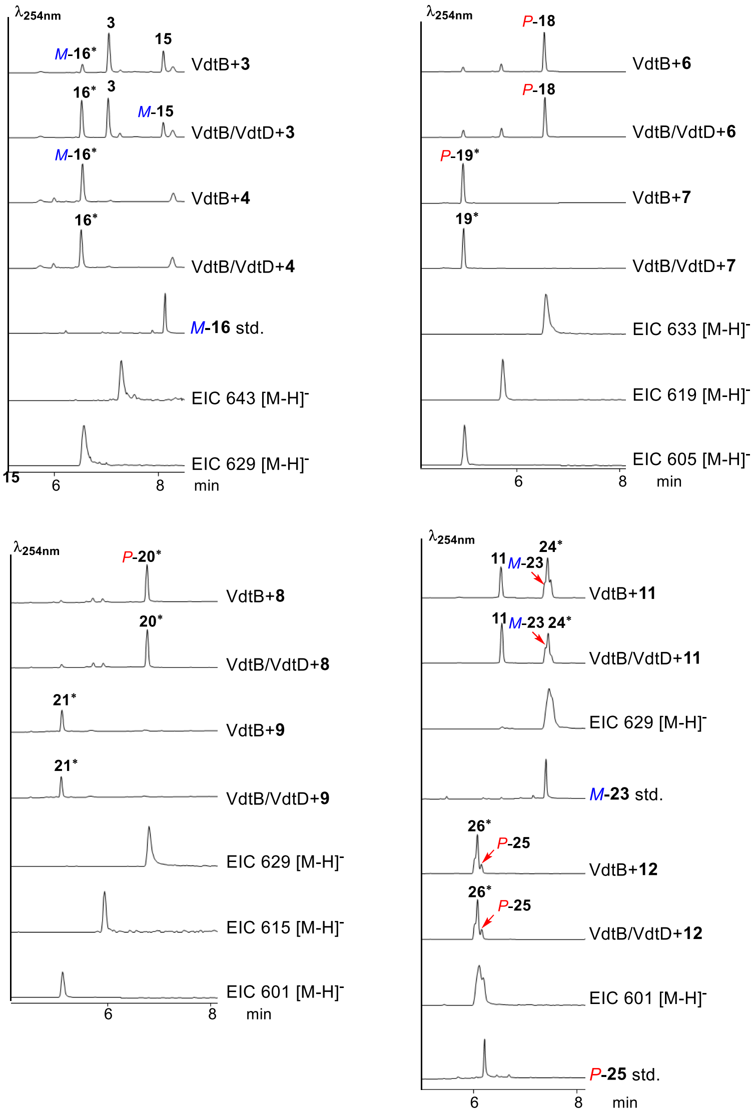

#### Figure S4. HPLC traces of heterologous expression of combinations mimicking the *in vitro* assay results

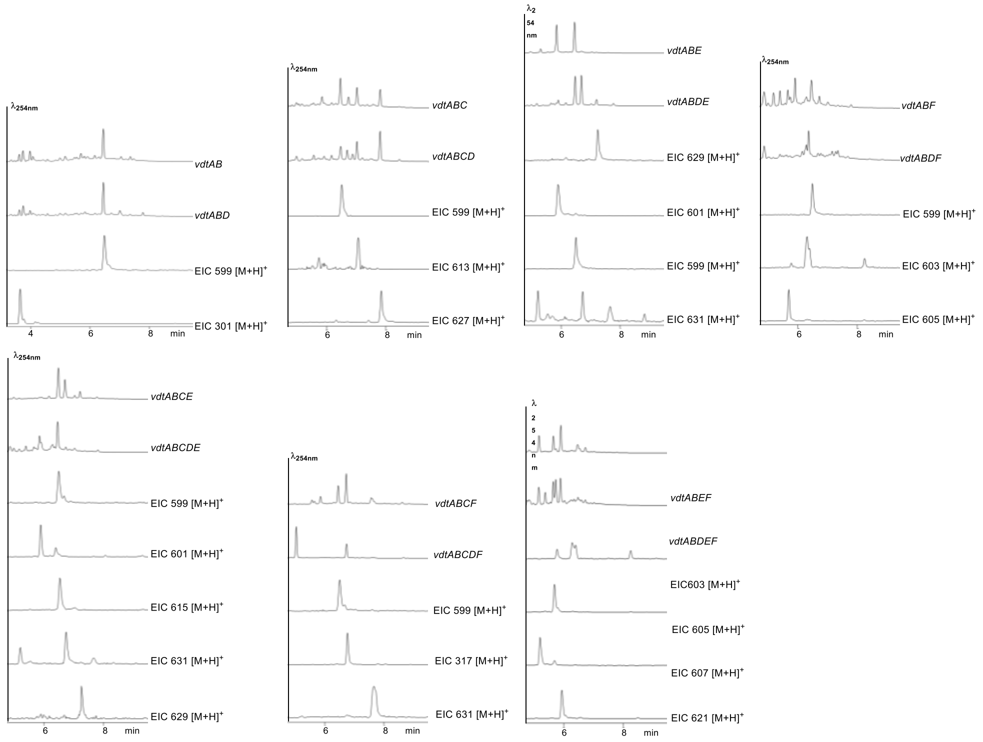

#### Figure S5. Characterized fungal BGCs encoding laccases/MCOs-mediated biaryl compounds biosynthesis.

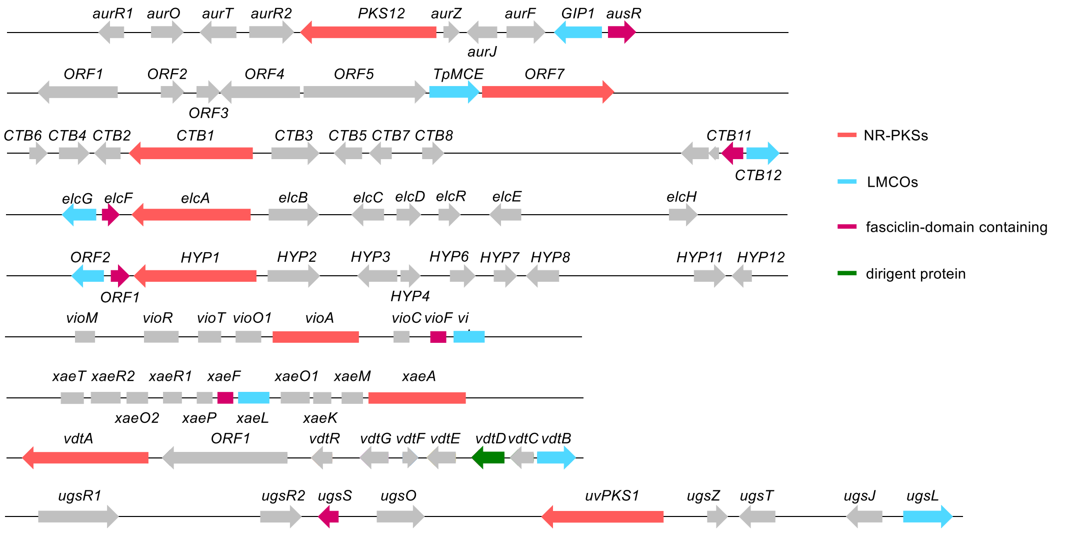

From top to bottom: aur cluster for biosynthesis of aurofusarin *Fusarium graminearum*;[4] *dpa* cluster for biosynthesis of dinapinones in *Talaromyces pinophilus*;[5] *elc*/*CTB*/*HYP* clusters for biosynthesis of elsinochrome C/cercosporin/hypocrellins;[1, 6] *vio* cluster for biosynthesis of vioxanthin in *Penicillium citreonigrum*;[7] *xae* cluster for biosynthesis of xanthoepocin in *Penicillium arizonense*;[7] *vdt* cluster for biosynthesis of viriditoxin in *Paecilomyces variotii*;[2] *ust* cluster for biosynthesis of ustilaginoidins in *Ustilaginoidea virens*.[8] The presence of fasciclin-domain containing proteins was highlighted.

#### Figure S6. *in vitro* assay of VdtD-D228S.

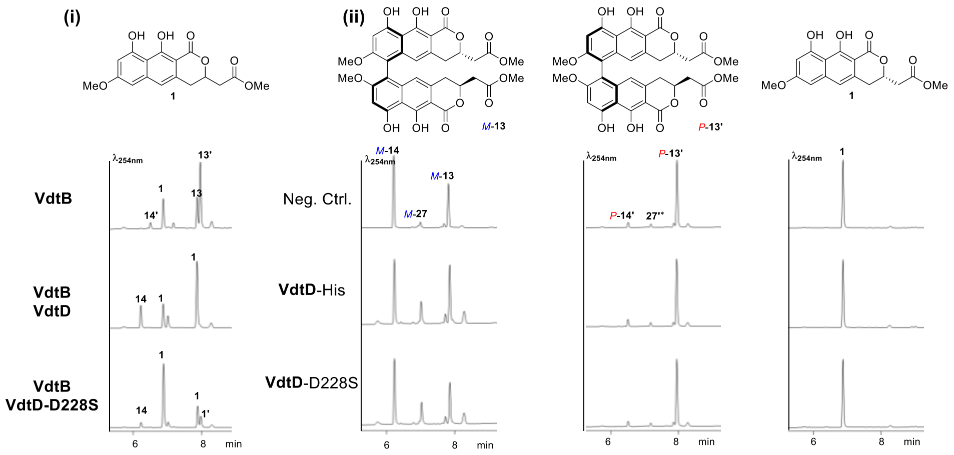

i) oxidative coupling of VdtB using **1** as substrate. VdtD-D228S still possesses stereoselective control over VdtB.

ii) **1**, **13**, and **13′** were used for testing the hydrolase activity of VdtD and VdtD-D228S. cell-free lysate of *A. nidulans* L0830 was used as negative control. It is noteworthy that the endogenous hydrolase act selectively on **13**. VdtD and VdtD-D228S, on the other hand, did not show obvious hydrolase activity.

#### Figure S7. *In vitro* assays of VdtB and VdtD with a mixture of 1 and 3 as substrates.

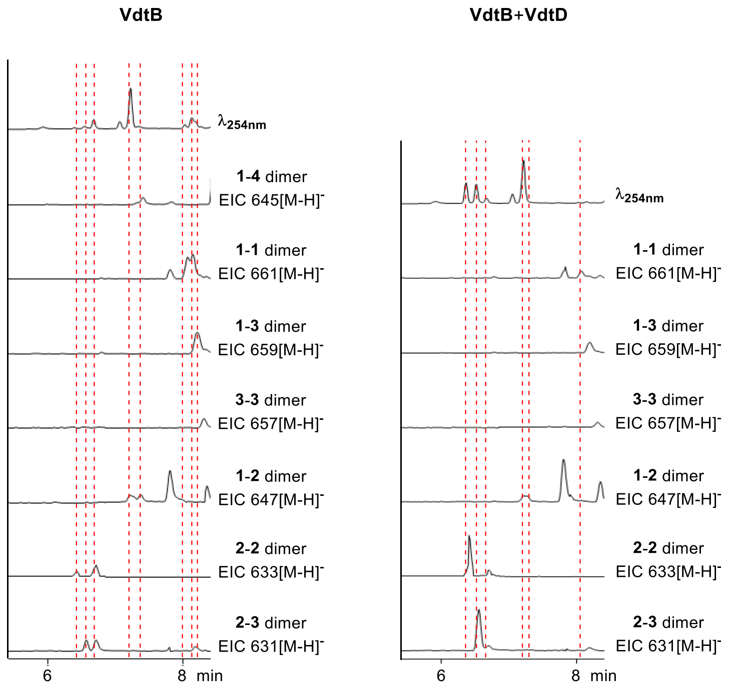

**Figure S8.** ***In vitro*** **assays of VdtB and VdtD with a mixture of 1 and 6 as substrates.**

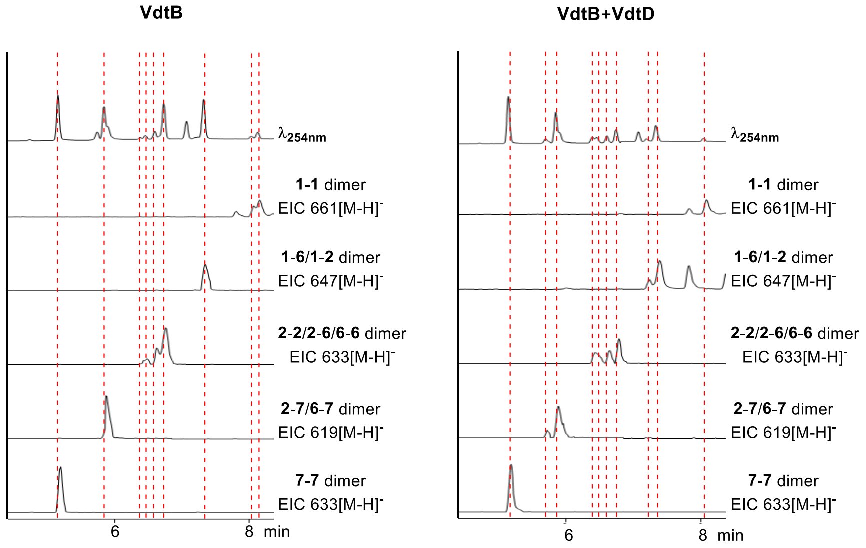

**Figure S9.** $^1$**H NMR spectrum (600 MHz) of 6 in DMSO-*d*$_4$**

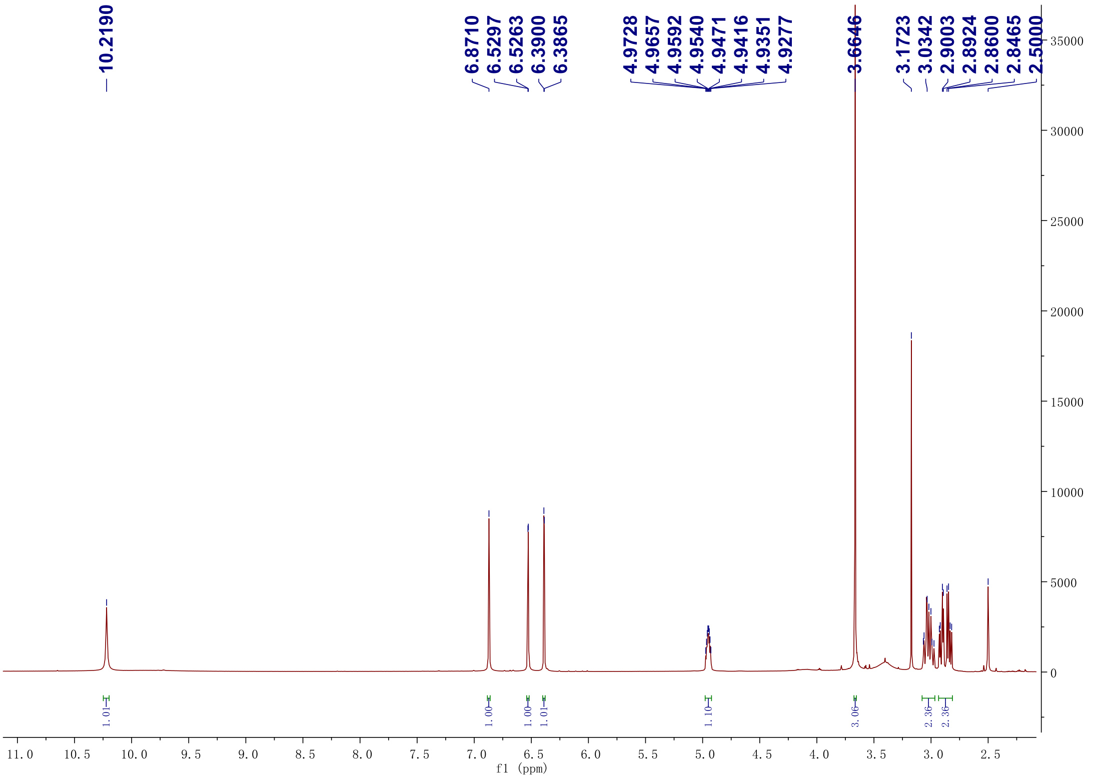

**Figure S10. $^{13}C$ NMR spectrum (150 MHz) of 6 in DMSO-$d_4$**

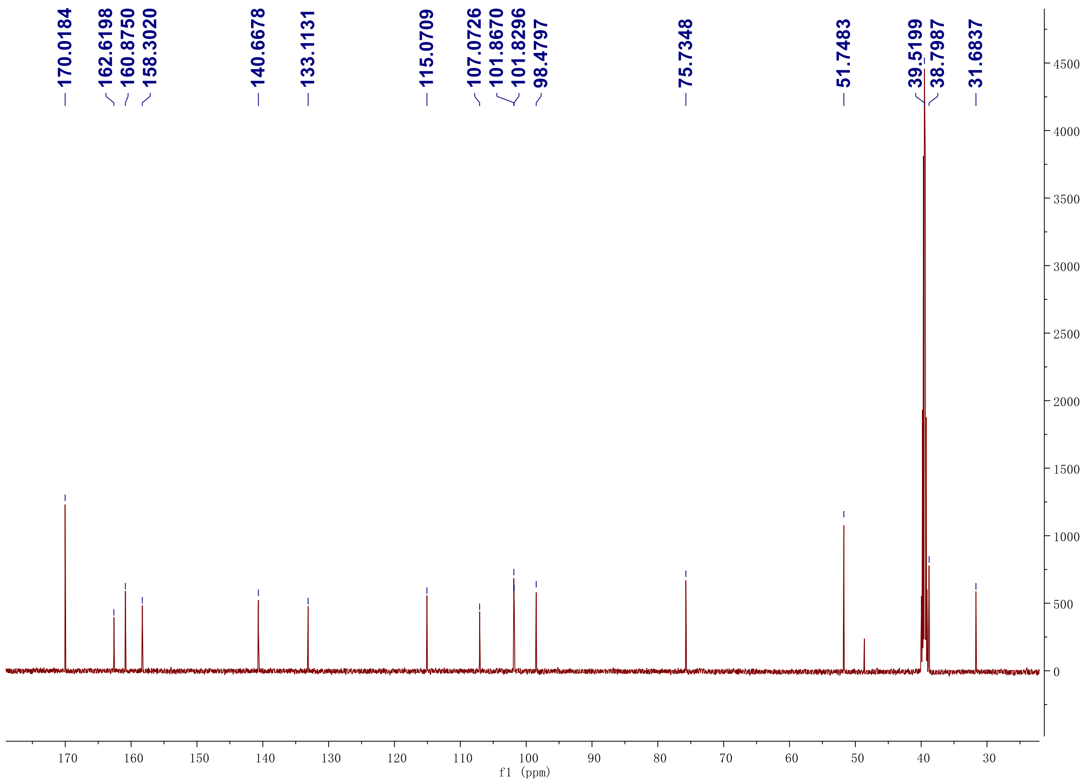

**Figure S11. $^{1}H$-$^{1}H$ gCOSY spectrum (600 MHz) of 6 in DMSO-$d_4$**

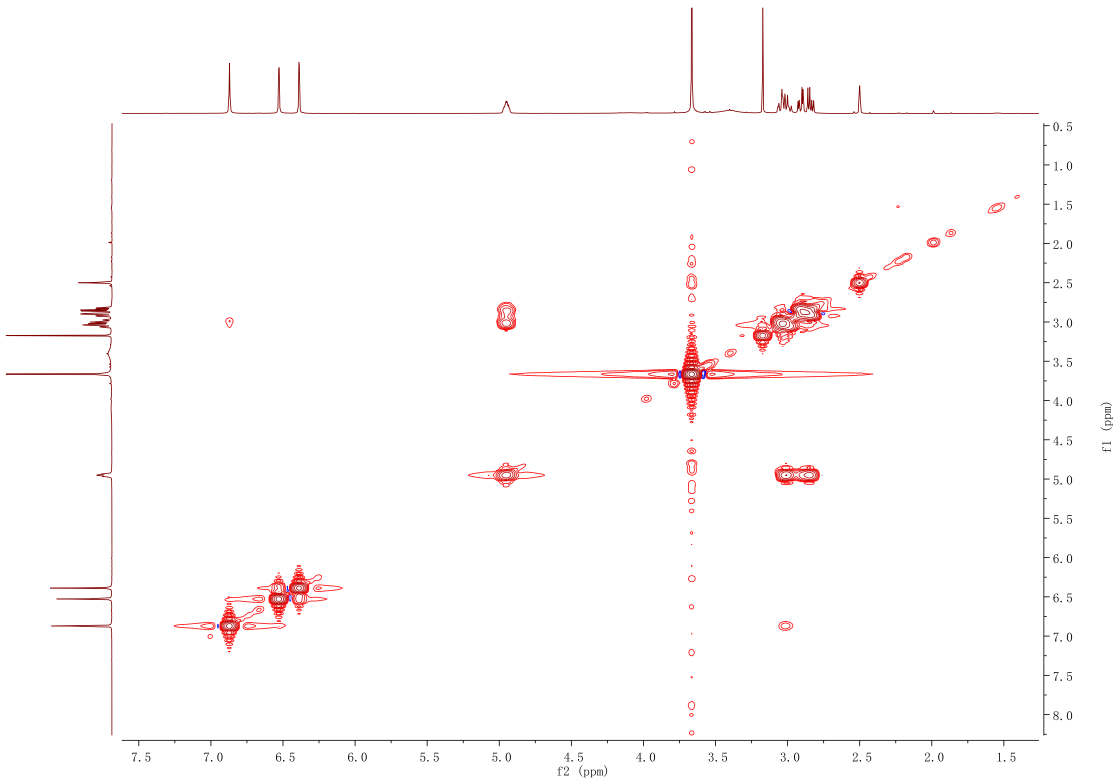

**Figure S12. HSQC NMR spectrum (600 MHz) of 6 in DMSO-$d_4$**

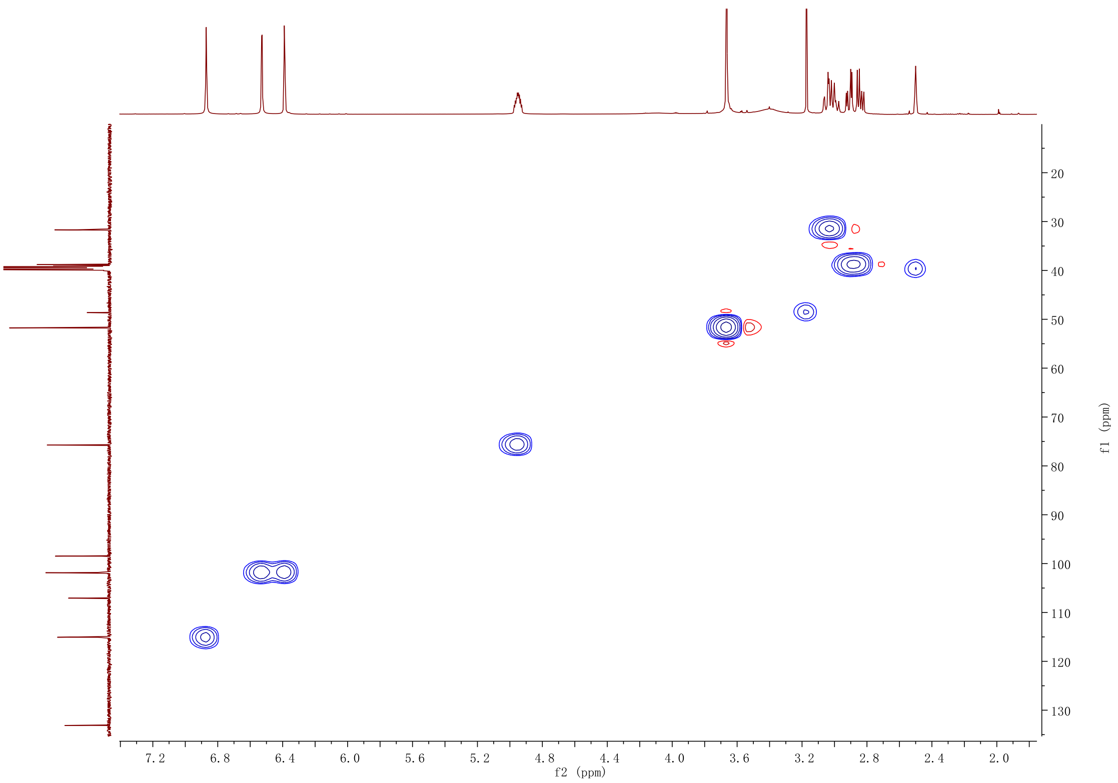

**Figure S13. HMBC NMR spectrum (600 MHz) of 6 in DMSO-$d_4$**

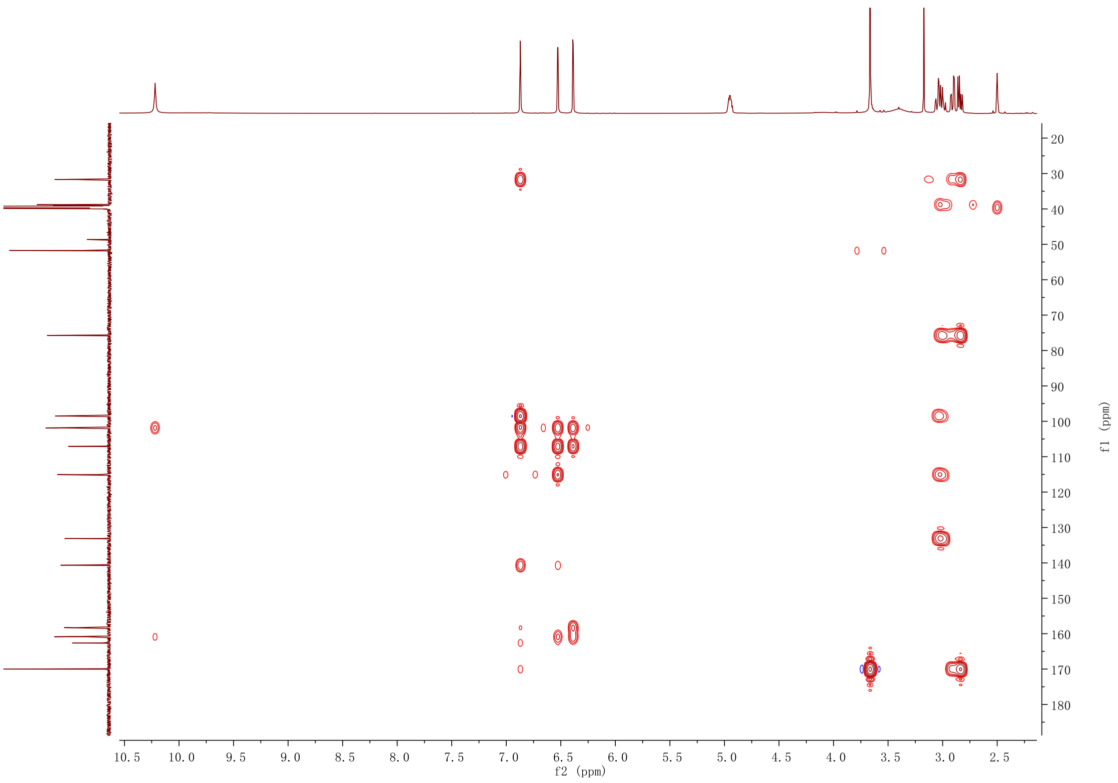

#### Figure S14. [1]H NMR spectrum (600 MHz) of 7 in methanol-$d_4$

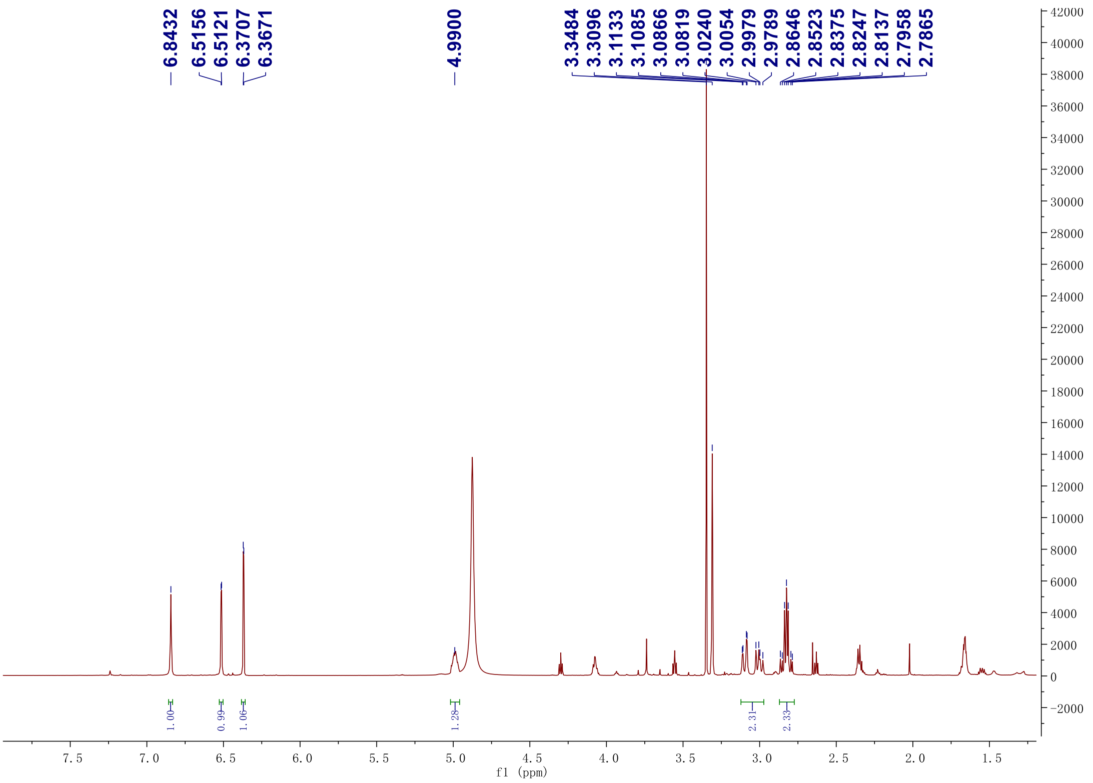

#### Figure S15. [13]C NMR spectrum (150 MHz) of 7 in methanol-$d_4$

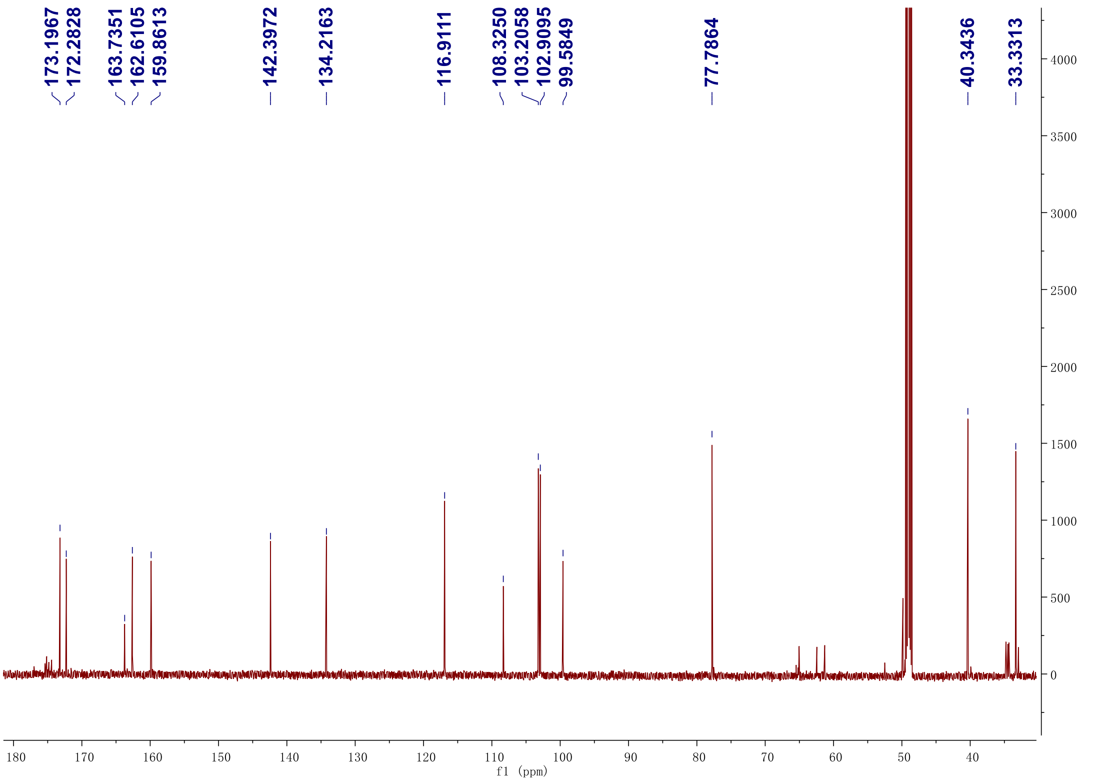

**Figure S16. $^1$H-$^1$H gCOSY spectrum (600 MHz) of 7 in methanol-$d_4$**

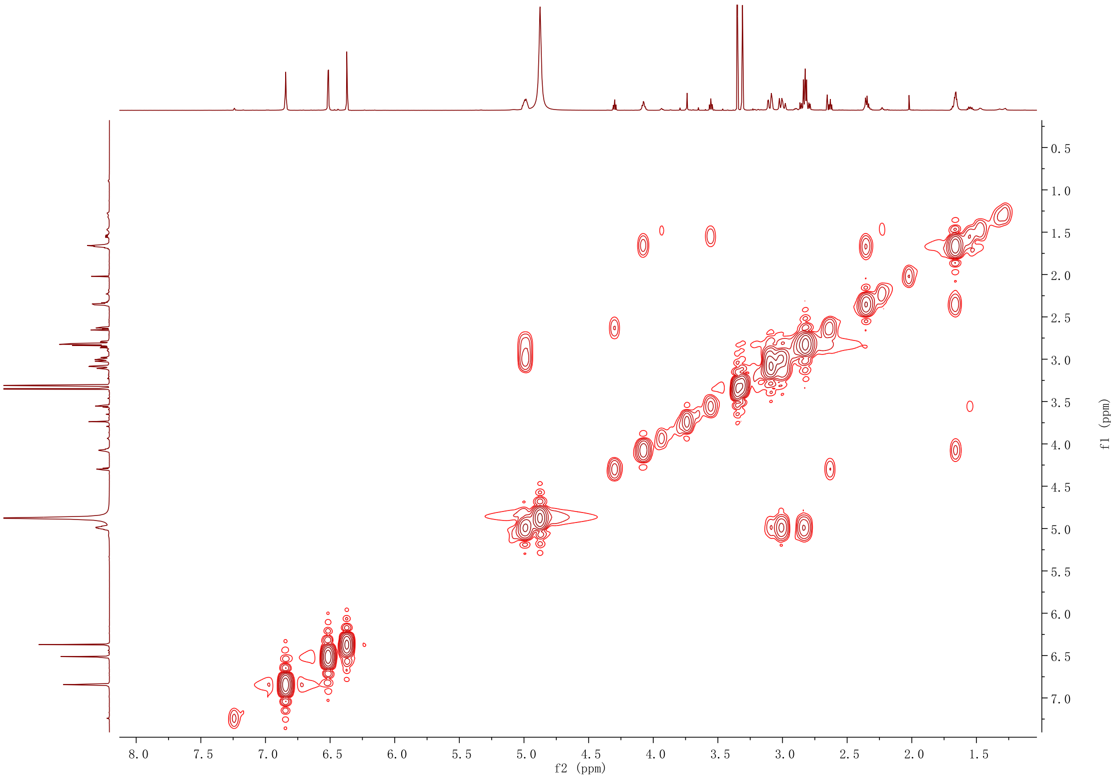

**Figure S17. HSQC NMR spectrum (600 MHz) of 7 in methanol-$d_4$**

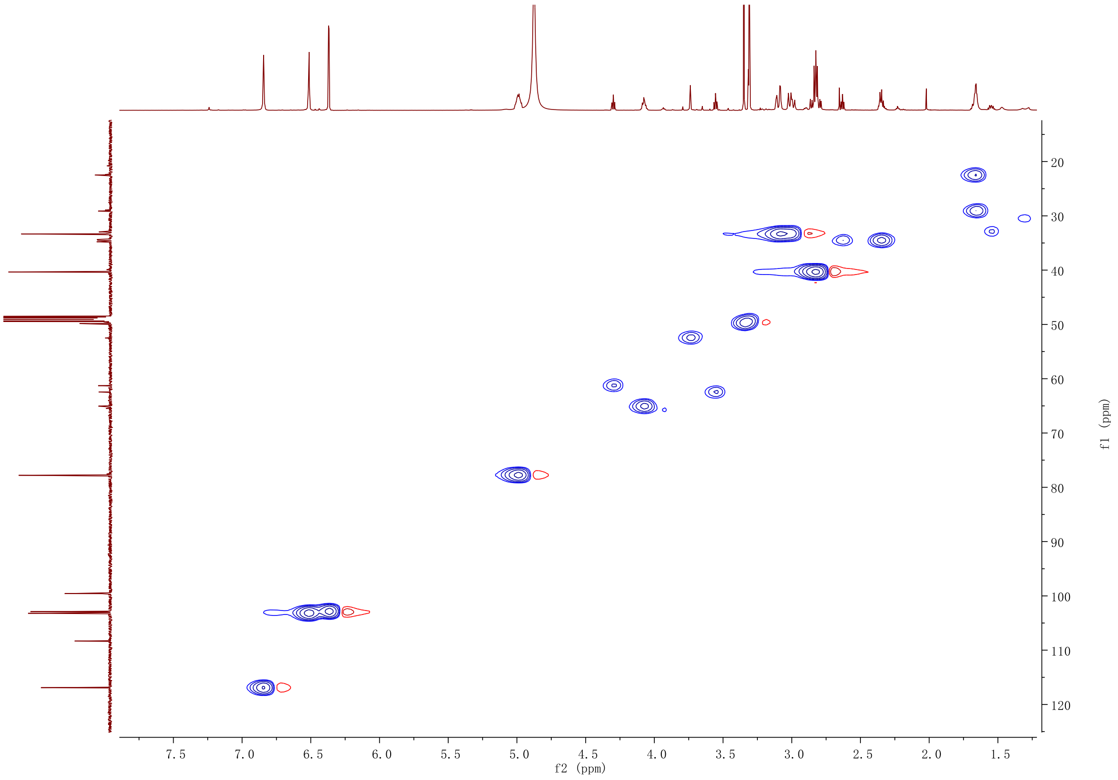

**Figure S18. HMBC NMR spectrum (600 MHz) of 7 in methanol-$d_4$**

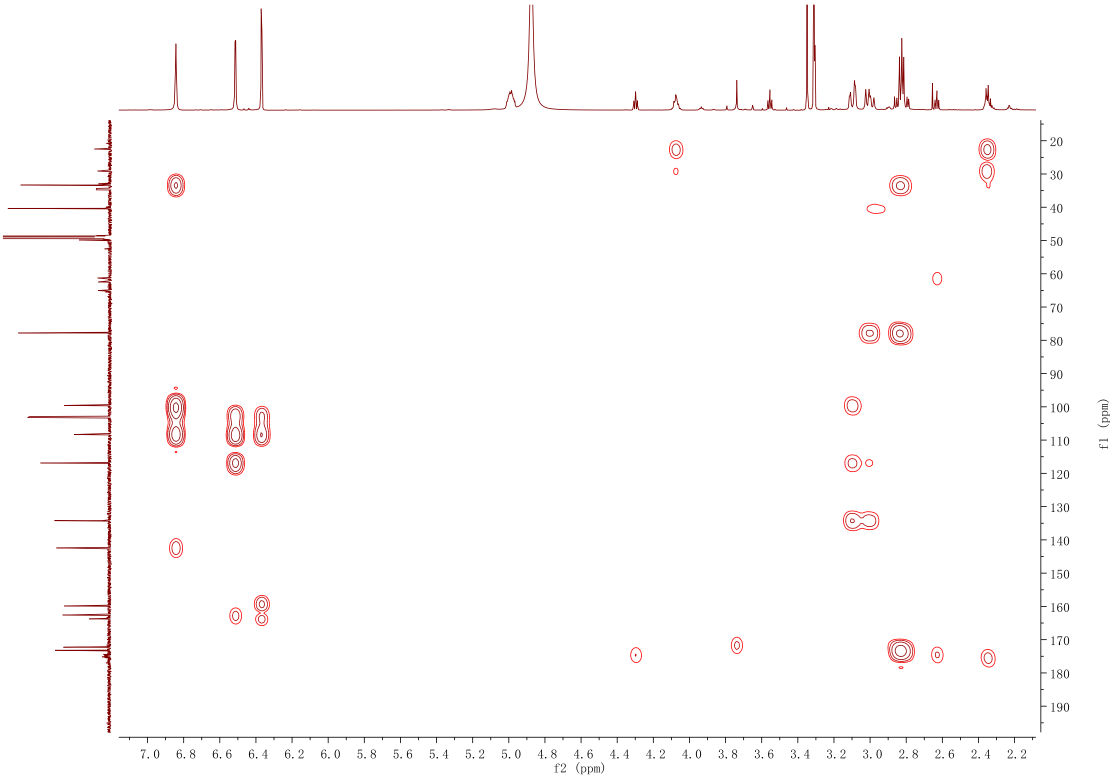

**Figure S19. $^1$H NMR spectrum (500 MHz) of 8 in DMSO-$d_4$**

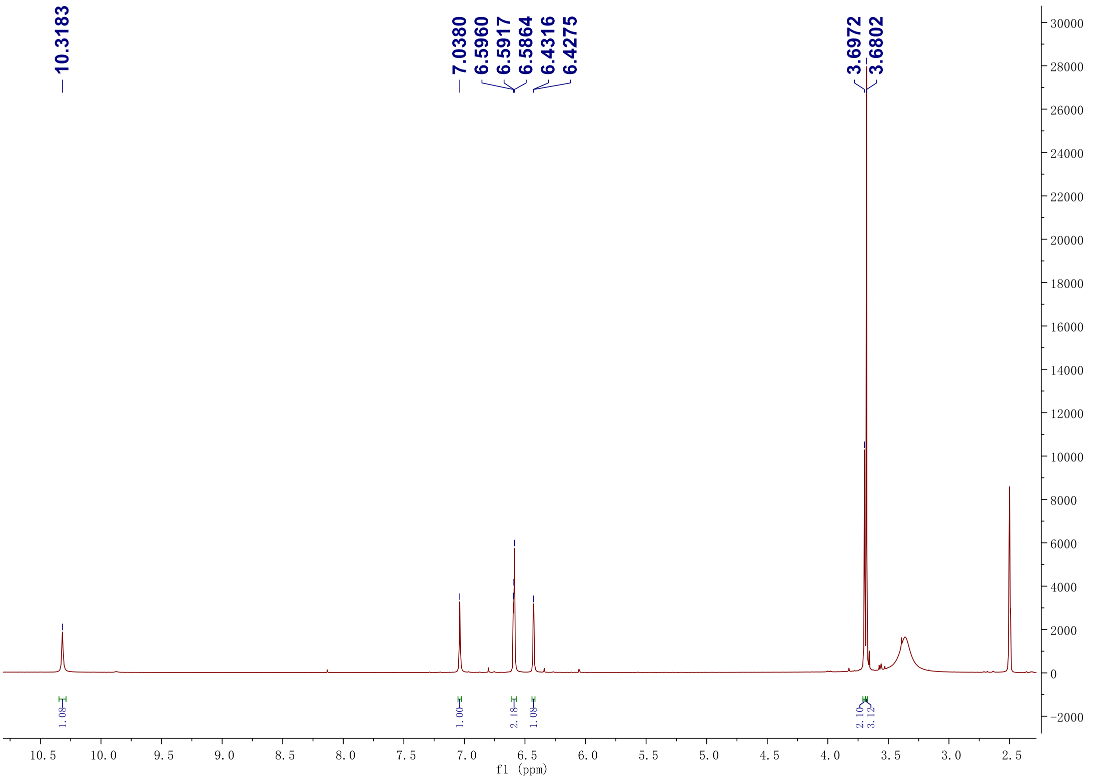

#### Figure S20. $^{13}$C NMR spectrum (125 MHz) of 8 in DMSO-$d_4$

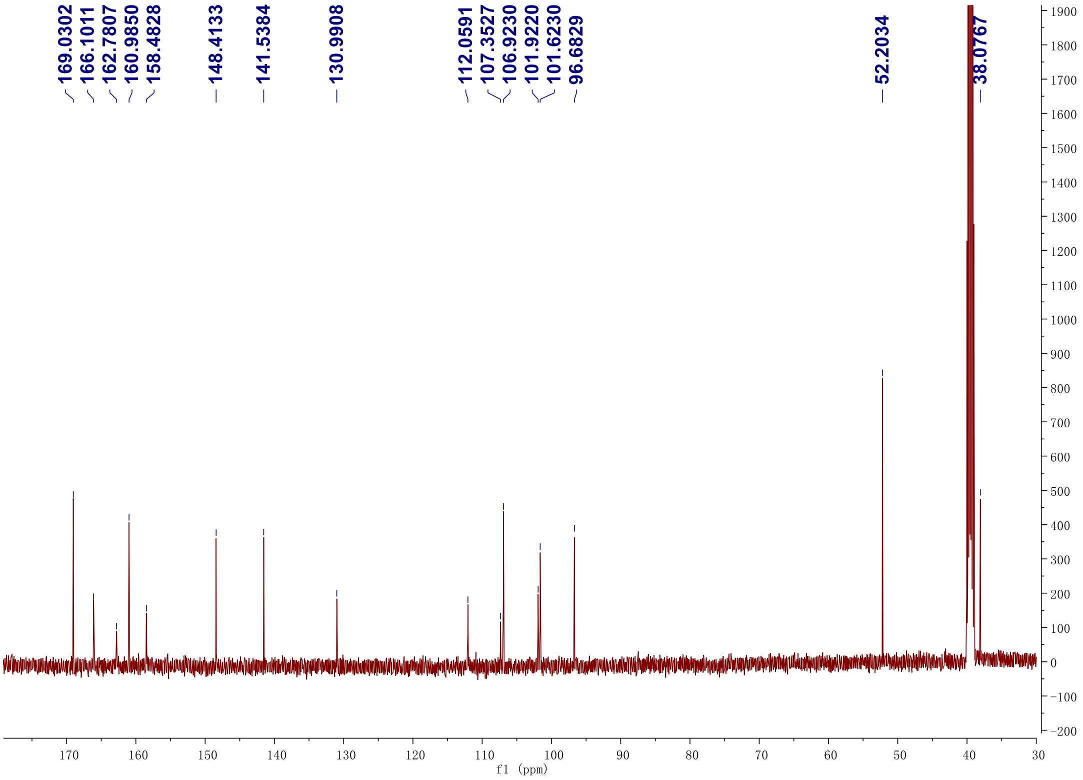

#### Figure S21. DEPT-135 $^{13}$C NMR spectrum (125 MHz) of 8 in DMSO-$d_4$

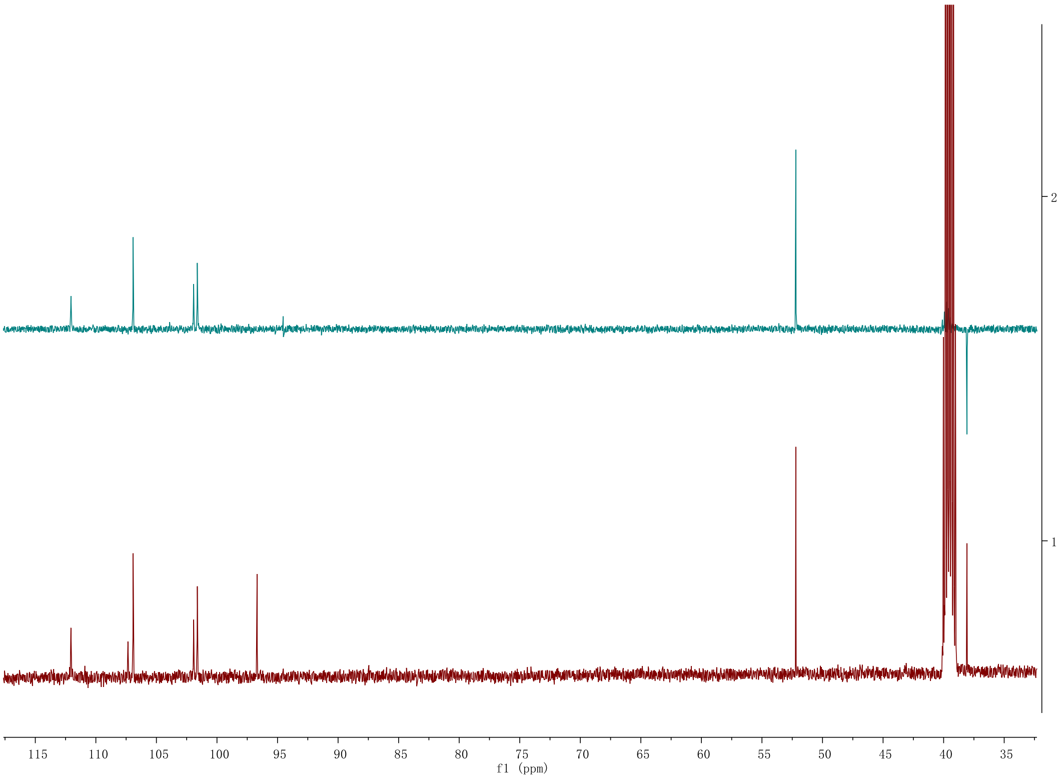

#### Figure S22. $^1H$-$^1H$ gCOSY spectrum (500 MHz) of 8 in DMSO-$d_4$

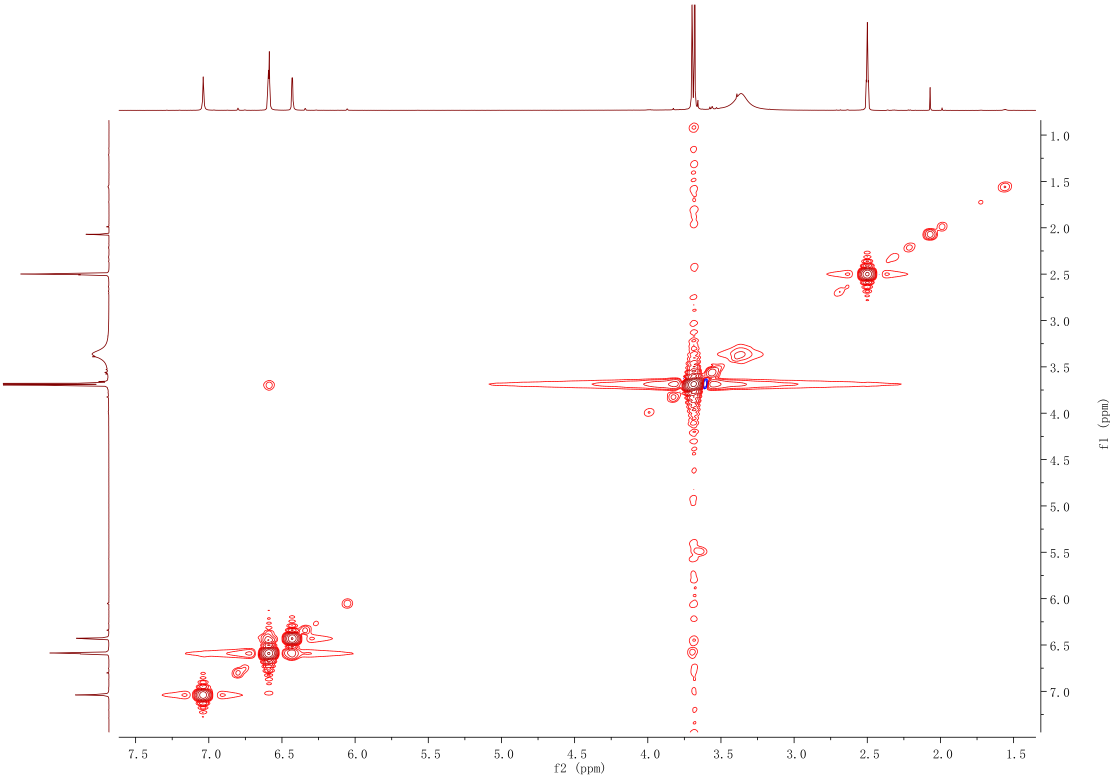

#### Figure S23. HSQC NMR spectrum (500 MHz) of 8 in DMSO-$d_4$

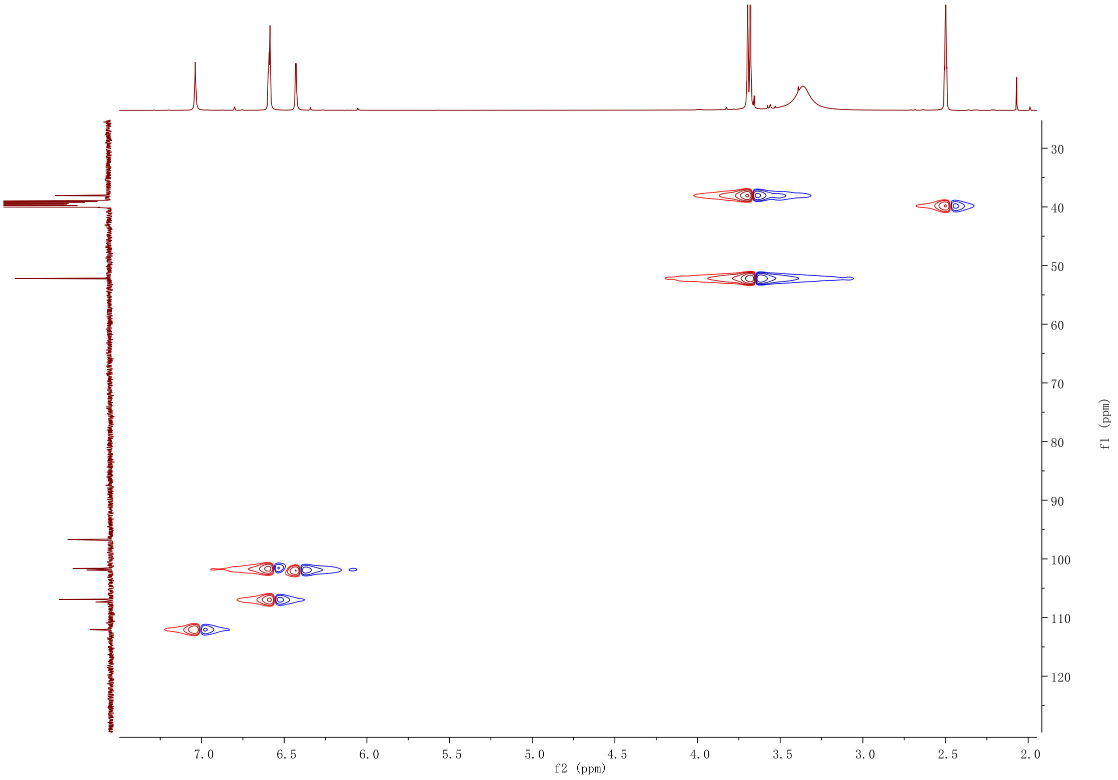

#### Figure S24. HMBC NMR spectrum (500 MHz) of 8 in DMSO-$d_4$

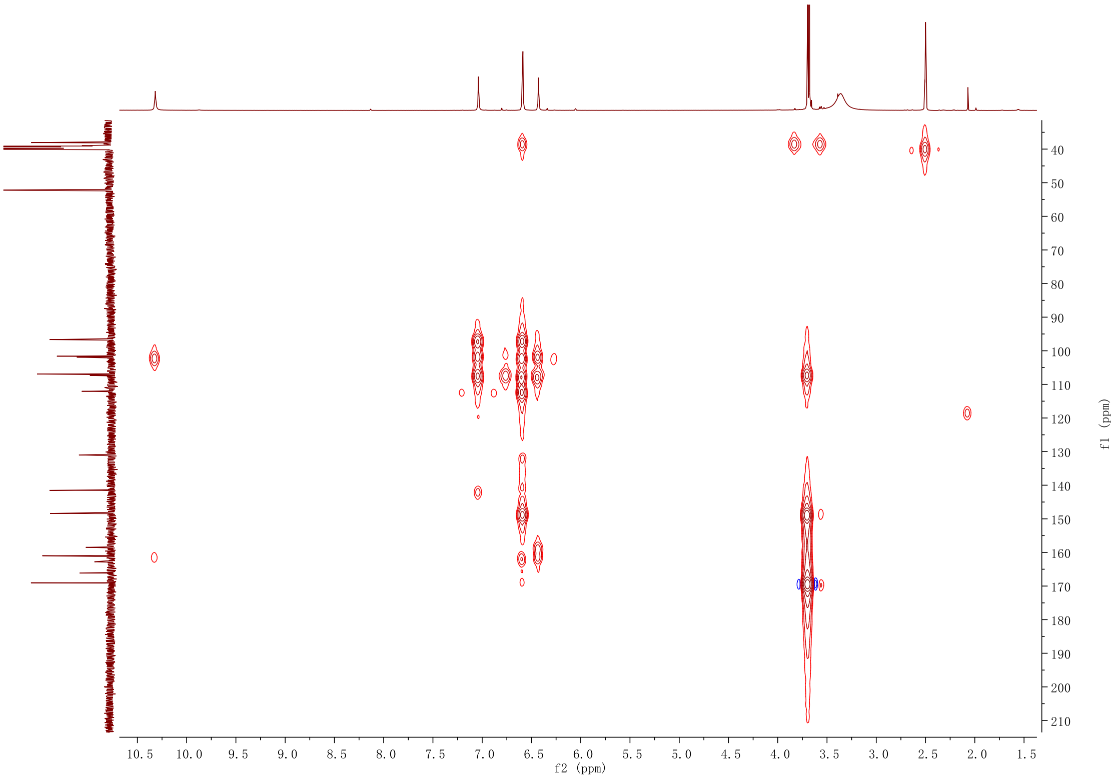

#### Figure S25. [1]H NMR spectrum (500 MHz) of 9 in DMSO-$d_4$

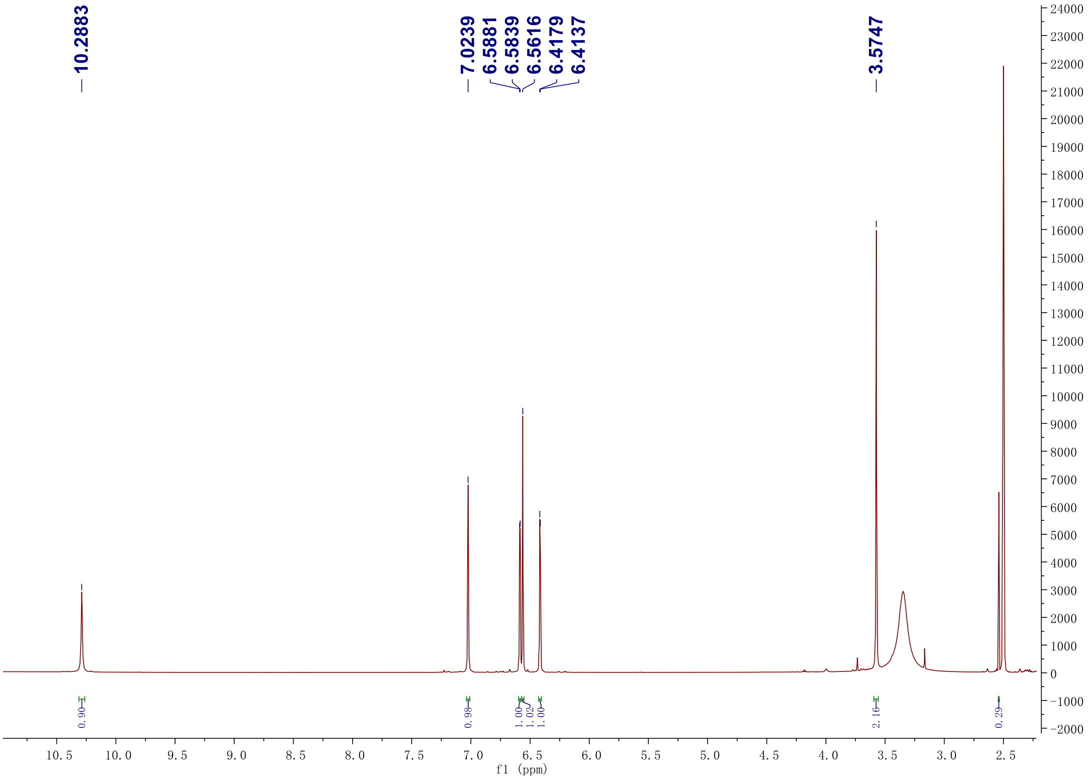

**Figure S26.** $^{13}C$ **NMR spectrum (125 MHz) of 9 in DMSO-$d_4$**

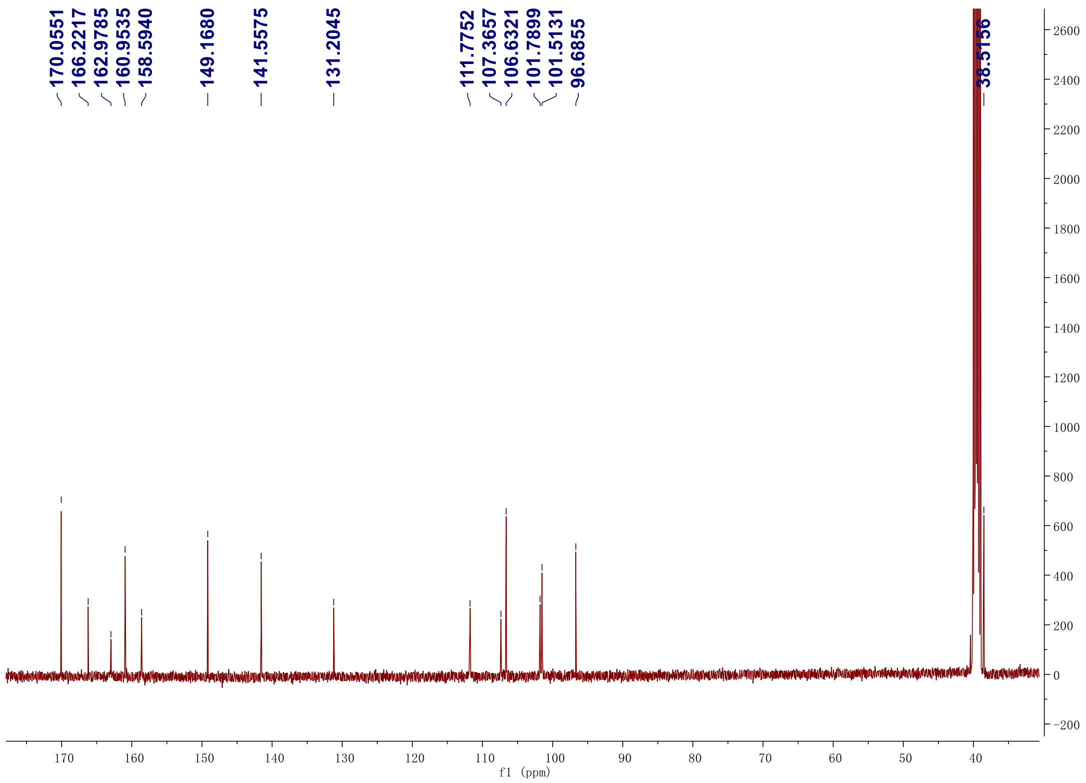

**Figure S27. DEPT-135** $^{13}C$ **NMR spectrum (125 MHz) of 9 in DMSO-$d_4$**

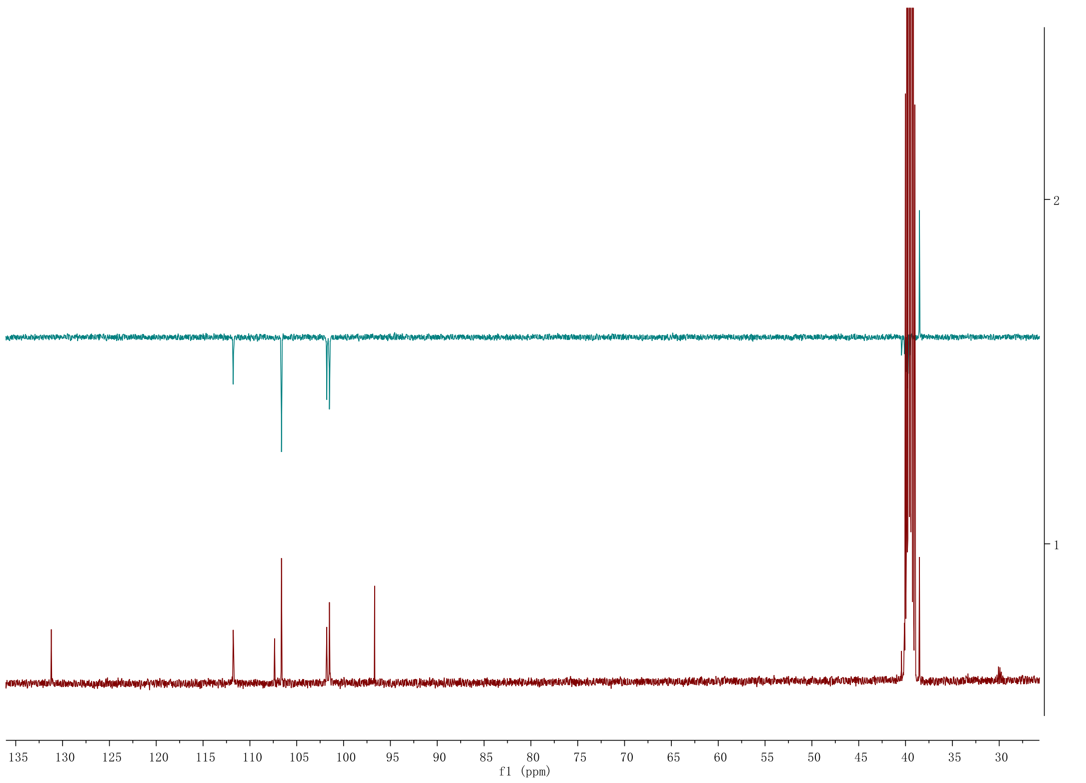

**Figure S28. [1]H-[1]H gCOSY spectrum (500 MHz) of 9 in DMSO-*d*4**

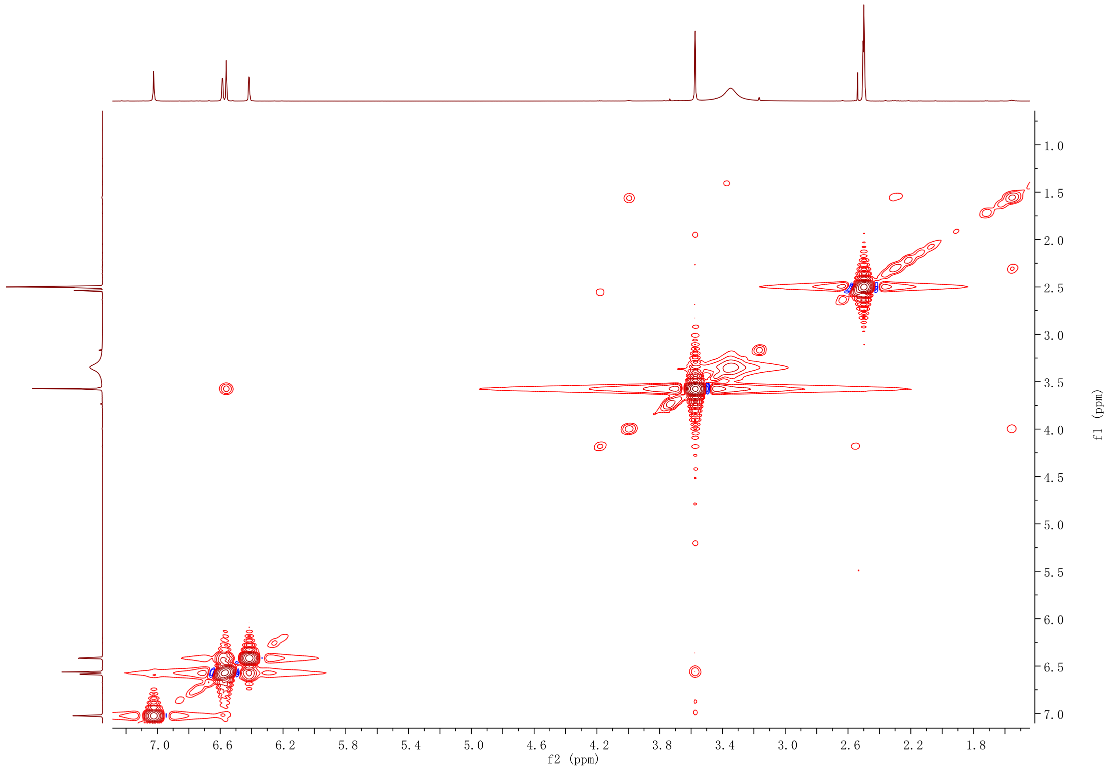

**Figure S29. HSQC NMR spectrum (500 MHz) of 9 in DMSO-*d*4**

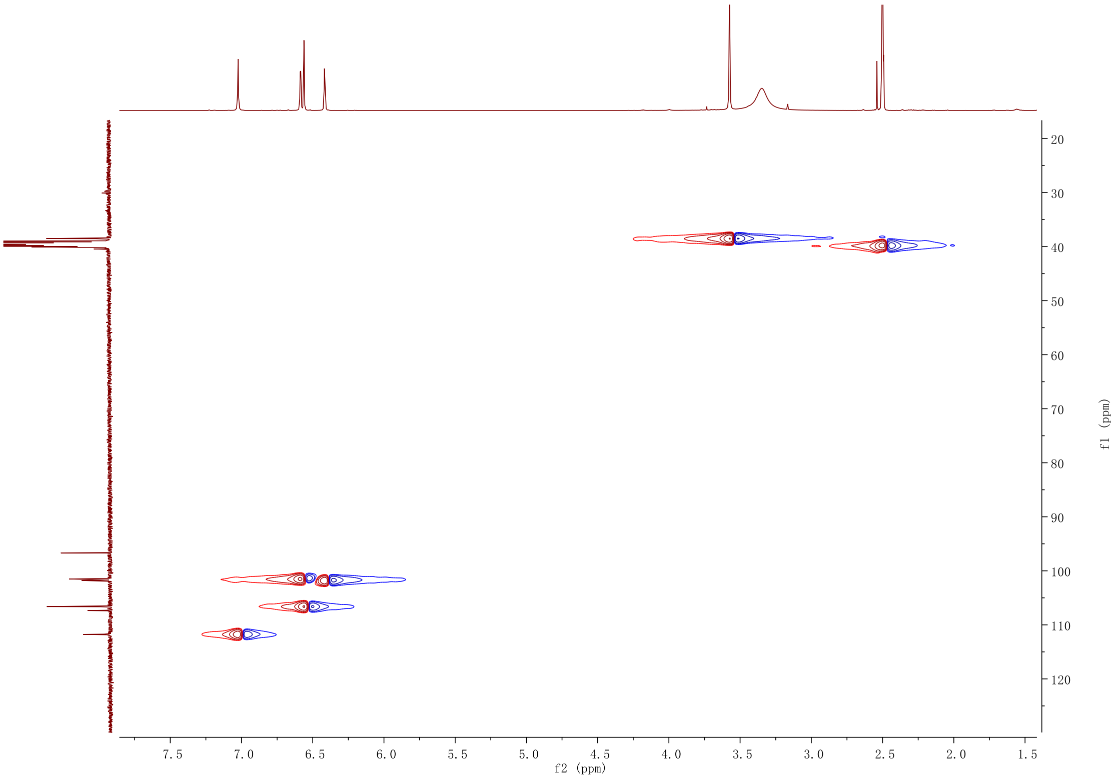

#### Figure S30. HMBC NMR spectrum (500 MHz) of 9 in DMSO-$d_4$

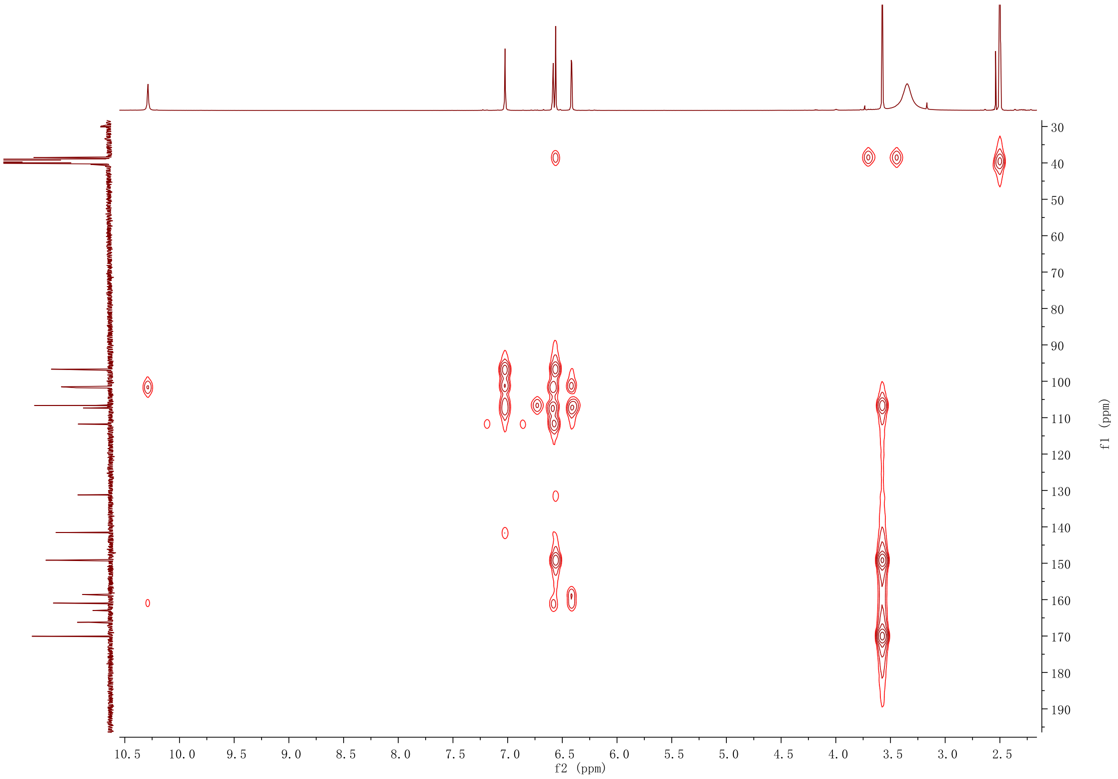

#### Figure S31. [1]H NMR spectrum (500 MHz) of 12 in DMSO-$d_4$

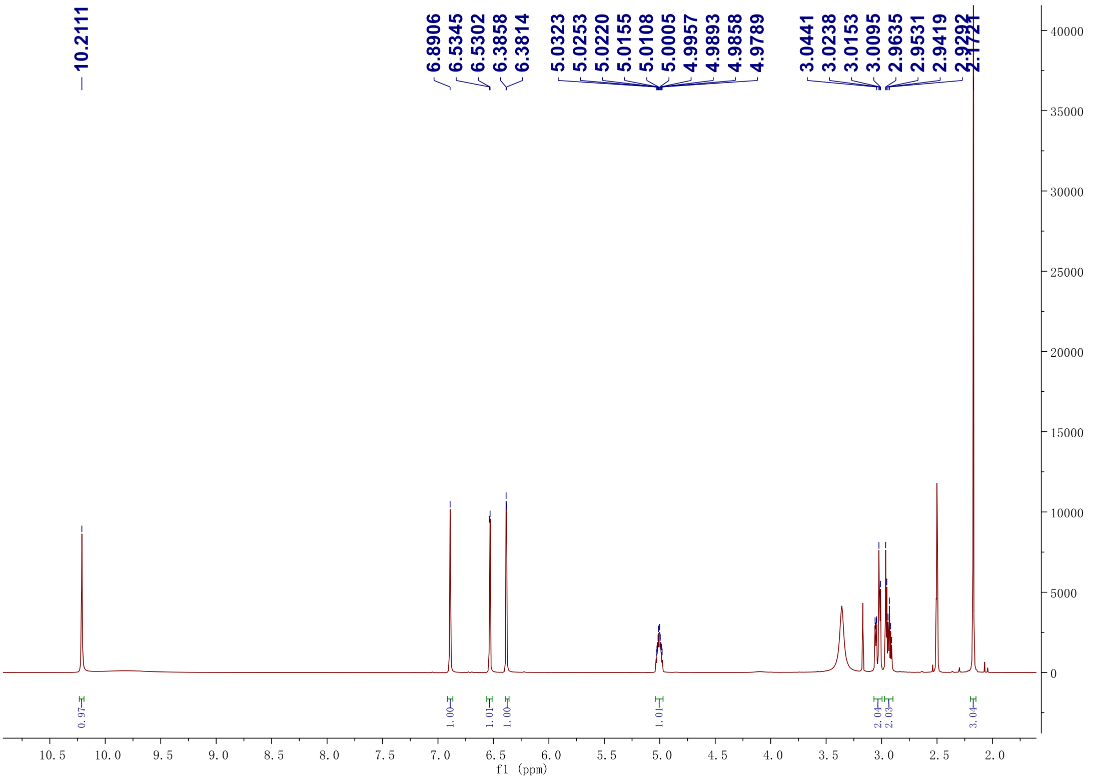

#### Figure S32. $^{13}C$ NMR spectrum (125 MHz) of 12 in DMSO-$d_4$

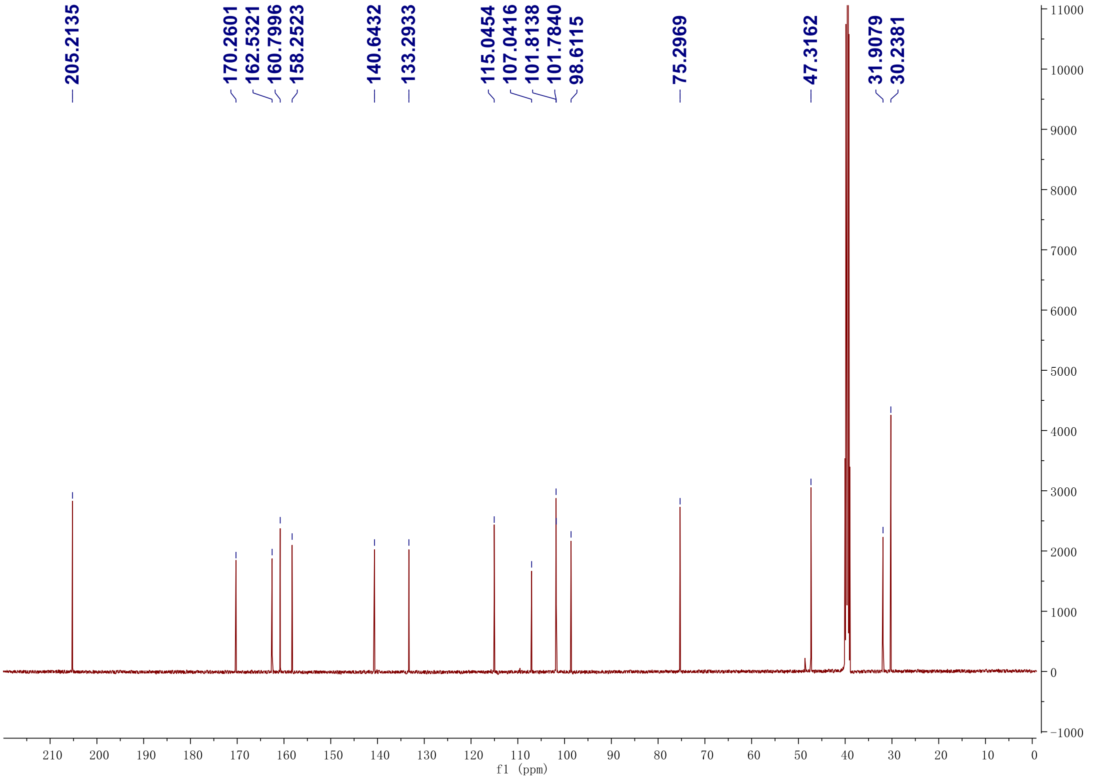

#### Figure S33. $^{1}H$-$^{1}H$ gCOSY spectrum (500 MHz) of 12 in DMSO-$d_4$

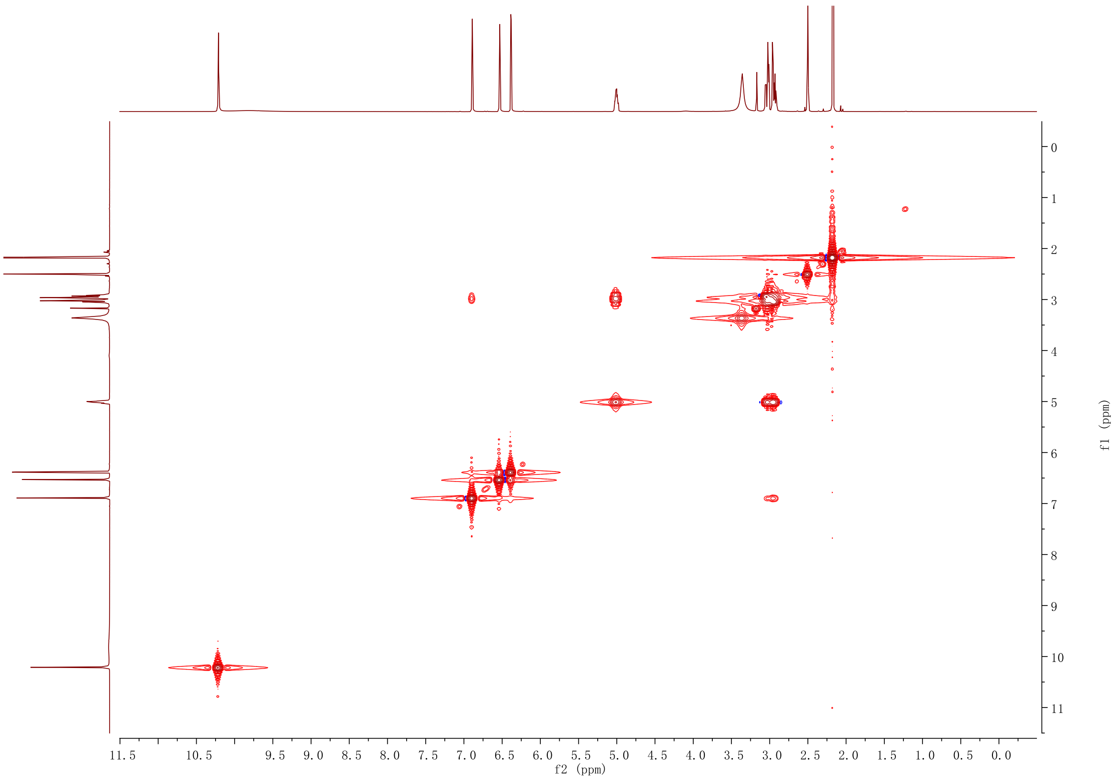

#### Figure S34. HSQC NMR spectrum (500 MHz) of 12 in DMSO-$d_4$

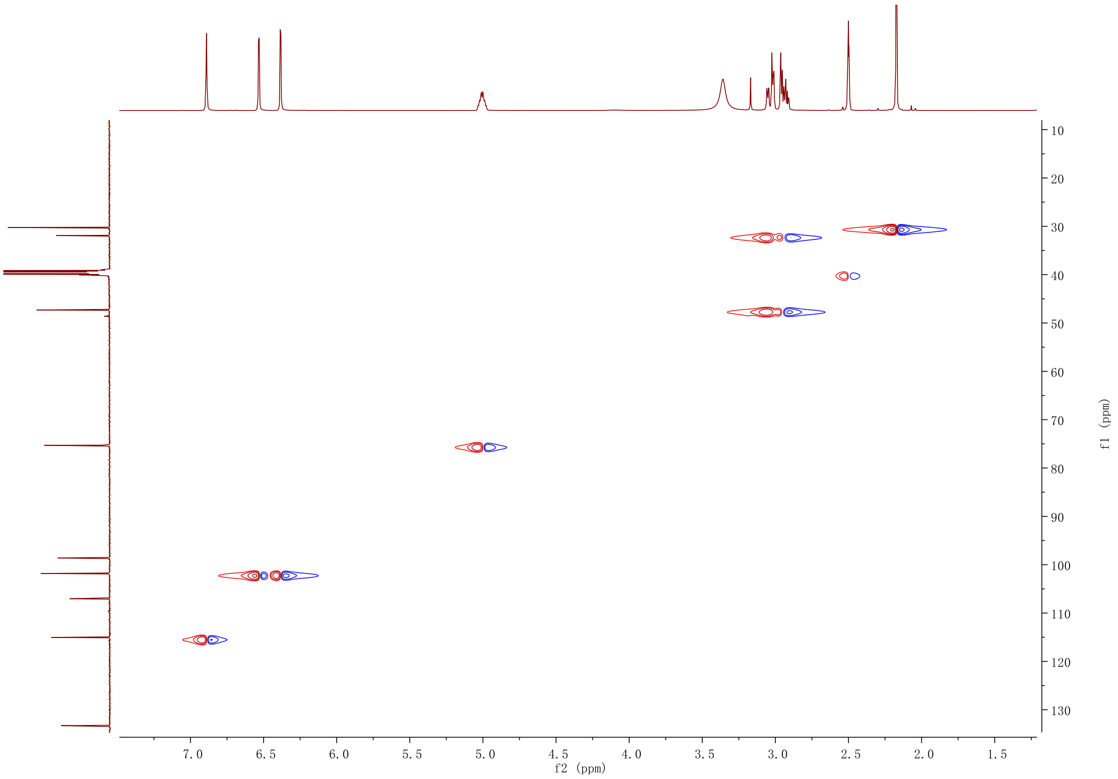

#### Figure S35. HMBC NMR spectrum (500 MHz) of 12 in DMSO-$d_4$

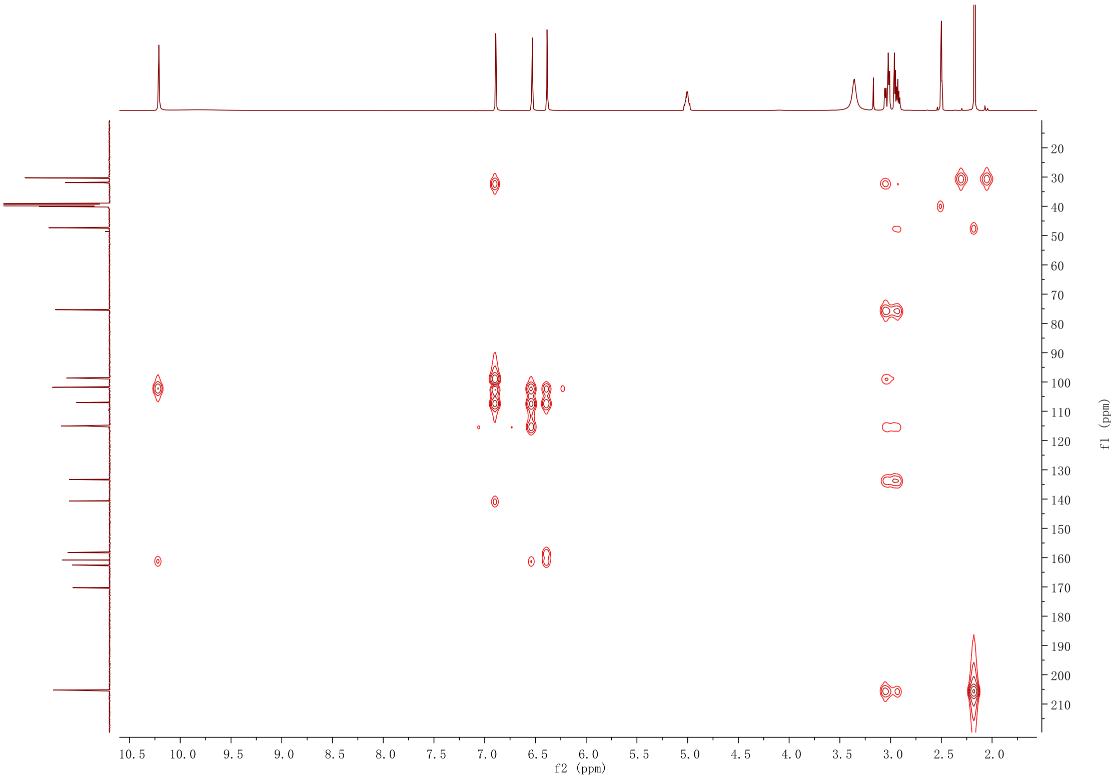

#### Figure S36. $^{1}H$ NMR spectrum (500 MHz) of 22 in acetone-$d_3$

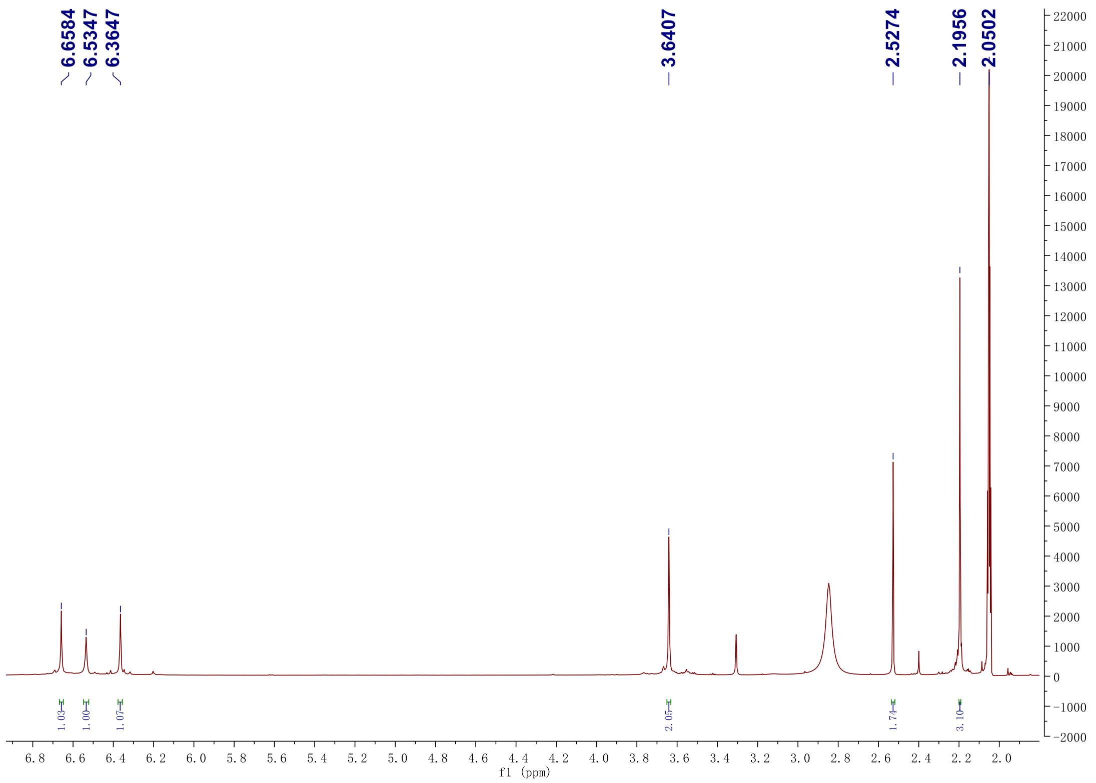

#### Figure S37. $^{13}C$ NMR spectrum (125 MHz) of 22 in acetone-$d_3$

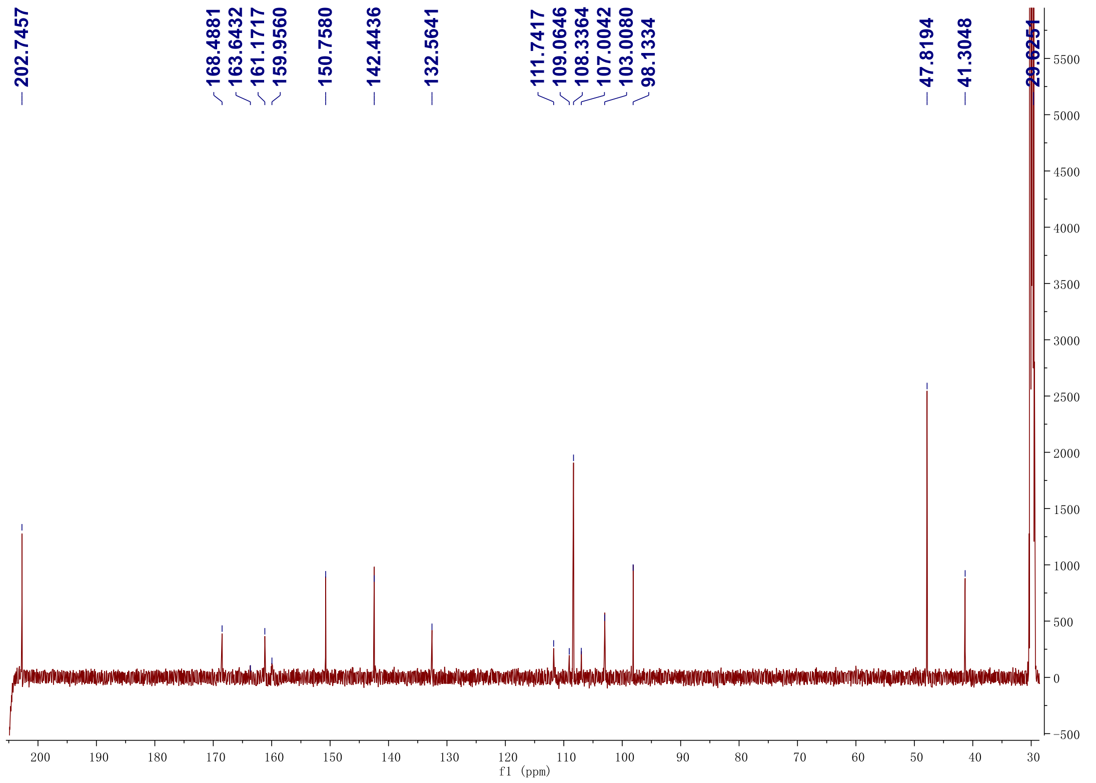

#### Figure S38. DEPT-135 $^{13}$C NMR spectrum (125 MHz) of 22 in acetone-$d_3$

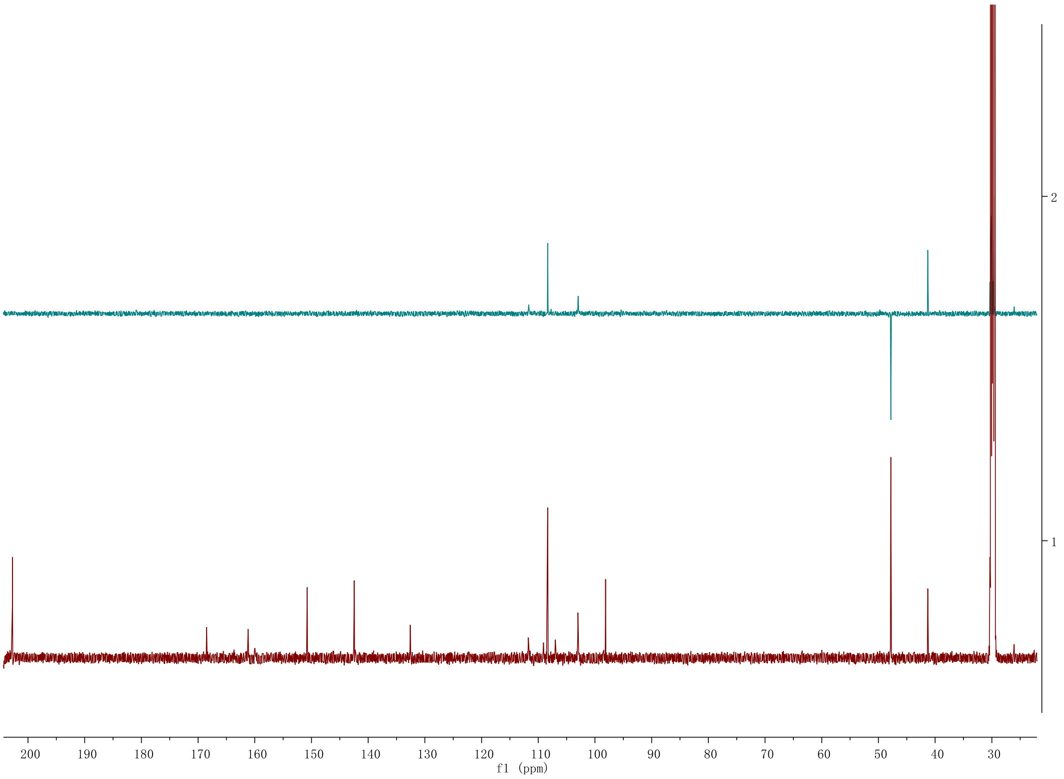

#### Figure S39. $^1$H-$^1$H gCOSY spectrum (500 MHz) of 22 in acetone-$d_3$

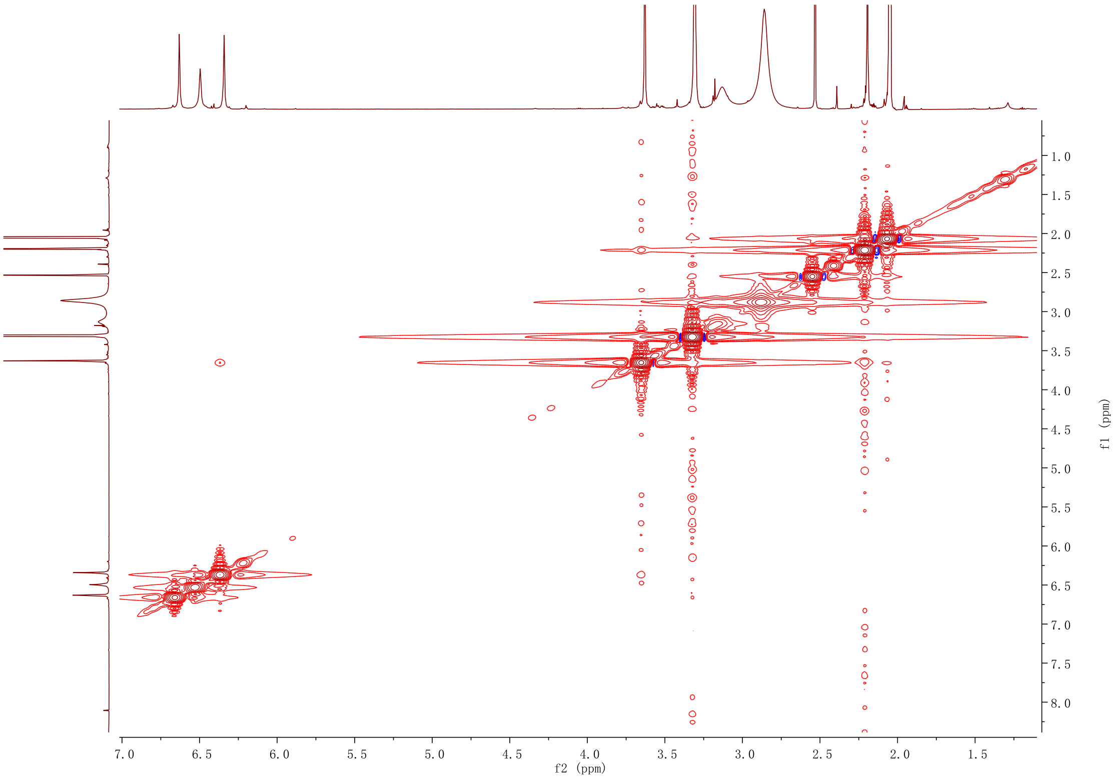

**Figure S40. HSQC NMR spectrum (500 MHz) of 22 in acetone-*d*₃**

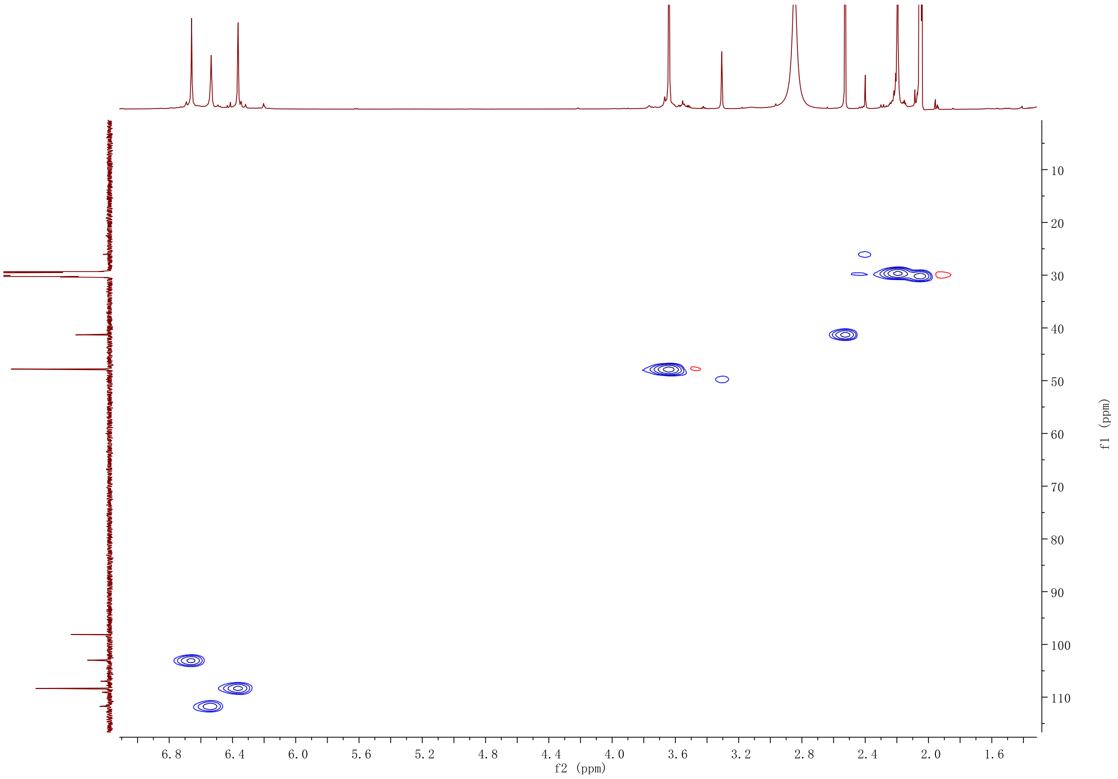

**Figure S41. HMBC NMR spectrum (500 MHz) of 22 in acetone-*d*₃**

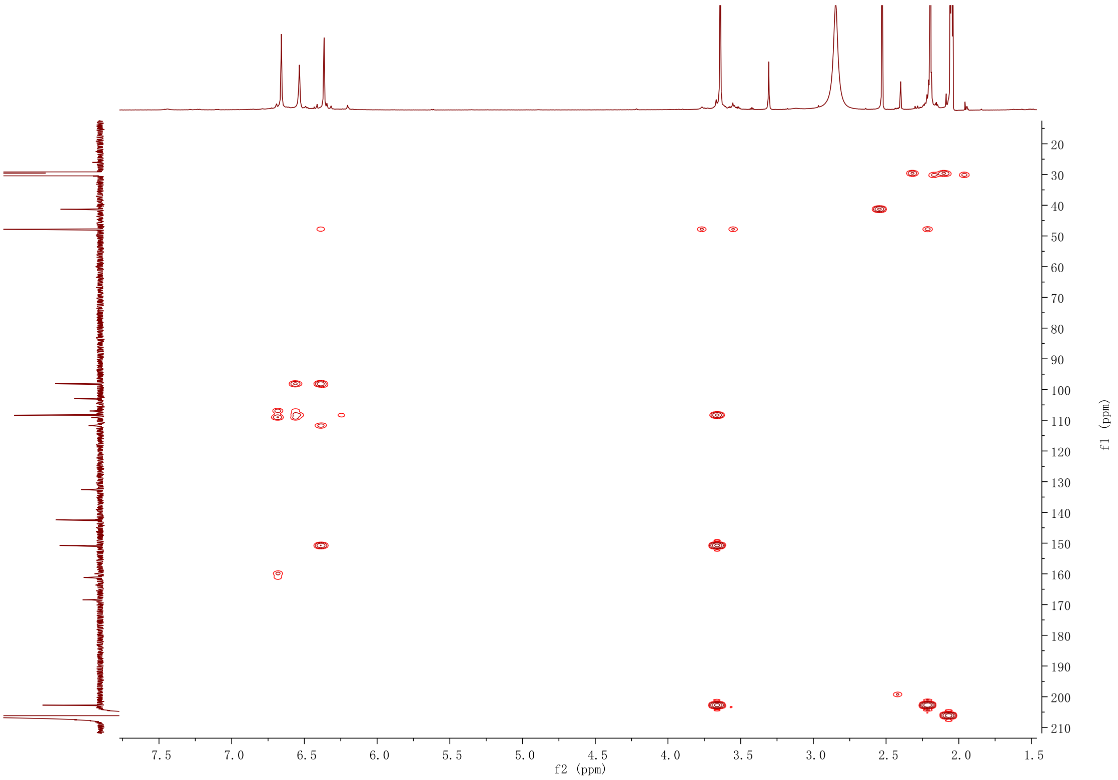

(6). De Jonge, R.; Ebert, M. K.; Huitt-Roehl, C. R.; Pal, P.; Suttle, J. C.; Spanner, R. E.; Neubauer, J. D.; Jurick, W. M.; Stott, K. A.; Secor, G. A., Gene cluster conservation provides insight into cercosporin biosynthesis and extends production to the genus *Colletotrichum*. *Proc. Natl. Acad. Sci. U. S. A.* **2018,** *115* (24), E5459-E5466.

(7). Fürtges, L.; Obermaier, S.; Thiele, W.; Foegen, S.; Müller, M., Diversity in Fungal Intermolecular Phenol Coupling of Polyketides–Regioselective Laccase-based systems. *ChemBioChem* **2019,** *20* (15), 1928-1932.

(8). Li, Y.; Wang, M.; Liu, Z.; Zhang, K.; Cui, F.; Sun, W., Towards understanding the biosynthetic pathway for ustilaginoidin mycotoxins in *Ustilaginoidea virens*. *Environ. Microbiol.* **2019,** *21* (8), 2629-2643.
